# Preconfigured cortico-thalamic neural dynamics constrain movement-associated thalamic activity

**DOI:** 10.1101/2023.09.20.558667

**Authors:** Perla González-Pereyra, Mario G. Martínez-Montalvo, Diana I. Ortega-Romero, Claudia I. Pérez-Díaz, Hugo Merchant, Luis A. Tellez, Pavel E. Rueda-Orozco

**Affiliations:** Departamento de Neurobiología del Desarrollo y Neurofisiología, Instituto de Neurobiología, UNAM; Departamento de Neurobiología Conductual y Cognitiva, Instituto de Neurobiología, UNAM

**Keywords:** Motor thalamus, Motor cortex, Preconfigured neural dynamics, Motor control

## Abstract

Neural preconfigured activity patterns (nPAPs) have been proposed as building blocks for cognitive and sensory processing. However, their existence and function in motor networks have not been explicitly studied. Here, we explore the possibility that nPAPs are present in the motor thalamus (VL/VM) and their potential contribution to motor-related activity. To this end, we developed a preparation where VL/VM multiunitary activity could be robustly recorded in mouse behavior evoked by primary motor cortex (M1) optogenetic stimulation and forelimb movements. VL/VM-evoked activity was organized as rigid stereotypical activity patterns at the single and population levels. These activity patterns were unable to dynamically adapt to different temporal architectures of M1 stimulation. Moreover, they were experience-independent and present in virtually all animals, confirming their preconfigured nature. Finally, subpopulations expressing specific M1-evoked patterns also displayed specific movement-related patterns. Our data demonstrate that the behaviorally related identity of specific neural subpopulations is tightly linked to nPAPs.

## Introduction

Neural activity patterns at the single and population levels have been proposed to underlie sensorimotor and cognitive processes. For example, memory formation in hippocampal networks has been associated with place cell sequential organization that can be recapitulated in different behavioral states such as environment exploration^1,2^, sleep^3^, or even short oscillatory periods^1,4^. On the other hand, in cortical and subcortical sensorimotor networks, neural population dynamics in the form of sequential activation have been associated with different aspects of movement, such as movement initiation^5^ or motor timing^6,7^. In turn, these complex population activity patterns appear to be composed of a variety of activity patterns at the single neuron level. For example, in the rodent cortical and subcortical areas, including input and output nuclei of the basal ganglia (BG), cutaneous whisker or forepaw stimulations induce diverse activity patterns at the individual level, generally characterized by a short-latency response, followed by a transitory inactivation period, and usually a second, long-latency response^8^**^‒^**^12^. This organization has also been reported for different sensory modalities, such as the auditory or olfactory systems^13^**^‒^**^15^, and is present in both behaving and anesthetized animals, suggesting a preconfigured nature. This stereotyped organization, sometimes referred to as “information packets”, spans for hundreds of milliseconds after stimulation and has been proposed as a generalized code for cognitive and sensory processing^2^. More recently, these neural preconfigured activity patterns (nPAPs) have been observed in the dorsolateral striatum in the context of motor timing^9^, where the experimental bidirectional manipulation of these dynamics has led to underestimations and overestimations of specific time intervals^9,16^.

While nPAPs have been linked to a variety of cognitive domains, including memory formation, sensory processing, and interval estimation, little is known about the implications of this apparently generalized code during movement production. In this context, previous reports have demonstrated that the motor cortical and thalamic areas (including the ventral lateral and ventral medial regions; VL/VM) are mutually indispensable to sustain movement-related activity^17^. On the other hand, previous observations have demonstrated that VL activity in primates and rodents efficiently predicts movement onsets hundreds of milliseconds in advance^17^**^‒^**^19^, and inactivation of VM delays movement onsets^20^. Furthermore, rhythmic auditory and visual stimulation evoke VL rhythmic neural patterns at single cell level, which can be used to timely predict movement onsets^21^. Finally, it has been proposed that a specific subgroup of pyramidal tract cortical neurons that projects to the motor thalamus is implicated in movement execution, but not preparation^22^, opening the possibility that VL/VM activity may also be segregated into different clusters related to specific movement parameters. Here we explored whether motor-related thalamic activity is constrained by preconfigured patterns of activity evoked by its main input, M1. First, we found that VL/VM activity evoked by general optogenetic M1 stimulation was organized into stereotypical activity patterns present in virtually all animals. These patterns were similar when slightly changing the temporal structure of the stimulation protocol, but most importantly, when comparing awake and anesthetized conditions and between naïve and highly trained animals in a forelimb movement task. Then, we analyzed if these M1-evoked VL/VM activity patterns were somehow related to specific behaviorally related neural activity patterns. By recording the same neurons during M1 stimulation and task performance, we found that VL/VM subpopulations expressing specific M1-evoked patterns also expressed specific movement-related patterns. Finally, optogenetic activation of the inhibitory pathway from the substantia nigra pars reticulata (SNr) to VL/VM regions disturbed nPAPs and movement execution parameters. Our data demonstrate that the behaviorally related identity of particular neural subpopulations is tightly linked to nPAPs.

## Results

### Motor thalamus responses to M1 stimulation are organized as stereotypical activity patterns

We performed silicon probe-based multiunitary activity recordings in the motor thalamus of head-fixed behaving mice (Fig. 1a). In the first set of experiments, recordings were performed in animals that were only accustomed to the head-fixed conditions for a period of 3 to 5 days; that is, no specific motor task was implemented. We targeted the M1–VL/VM motor thalamus circuit. We expressed and stimulated channelrhodopsin 2 (ChR2) in M1 and recorded neural dynamics in VL/VM regions (Fig. 1b-c). This general manipulation would impact on multiple groups of M1 projecting neurons, including intratelencephalic and cortico-cortical neurons not projecting to the thalamus, and pyramidal tract and cortico-thalamic neurons, known to project to VL/VM^23^. Based on previous evidence from cortical and subcortical networks ^9,11,24^, we first explored the possibility that VL/VM activity would reflect passive cortical stimulation (i.e., non-associated with specific behaviors or events) as repeatable neural dynamics at the single and population levels. To this end, we optogenetically stimulated M1 with two different trains of five stimuli differing in inter-stimulus intervals (ISIs; 300 ms, 3.3 Hz and 500 ms, 2 Hz). We chose the range of 300 - 500 ms ISI based on previous reports indicating this frequency as a proxy of the speed of forelimb movements in rodents when walking or trotting^25–27^. In this set of experiments, we recorded a total of 1018 neurons from 18 awake mice (1 to 3 recording sessions per animal). From these neurons, 641 presented stable responses to at least one stimulation protocol and from those, 377 presented stable responses in both stimulation protocols (see methods). The stimulation protocols produced a variety of activation patterns easily visible in the raster plots (Fig. 1d). As a population, neurons presented baseline firing rates consistent with previous reports (median 3 Hz, 25^th^ and 75^th^ percentiles, 1.3 to 5.5 Hz; Fig. 1e). A general visual examination of the neural activity suggested the M1-evoked responses were characterized by short-latency transient activation, followed by a brief period of inactivation, and a long-latency rebound (Fig. 1f). Most neurons appeared to maintain their response pattern in both stimulation protocols (300 and 500 ms ISI), but to formally explore a potential adaptation to ISIs, we first quantified the response latency for the three main components of the response. Histograms confirmed the presence of two groups of latencies for the incremental responses and one group for the decremental responses at the population level and for both ISIs (Fig. 1g-h). When comparing these latencies between ISIs, we found that the short-latency responses (increase and decrease) were similar. However, for the long-latency increases, the 500 ms ISI presented significantly shorter latencies than the 300 ms ISI (Fig. 1i). The median short-latency response was about 10 ms for ISIs, suggesting this is the direct M1-VL/VM connection. For their part, the decrease and rebound responses may be related to intrinsic properties of the cells or micro/macro circuit mechanisms. Then we analyzed if the magnitude of the M1-evoked responses could experience short-term adaptation to the different ISIs^28^. First, the average response of all neurons confirmed that the prototypical pattern of increase-decrease-increase in spiking activity was maintained for each stimulus of the train and in both ISIs (Fig. 1j). However, when we compared the amplitude of the short-latency increase, we observed that even when both conditions presented similar average trends, only for the 300 ms ISI, the second to fifth stimulus of the train were significantly larger when compared with the first stimulus (Fig. 1k), suggesting a mechanism for short-term facilitation depending on the frequency of stimulation in the M1-VL/VM connection. The amplitude of the decrease response was not significantly different between stimuli or ISI conditions (Fig. 1l). Finally, for both ISIs, the last two stimuli of the train produced significantly smaller amplitudes when compared with the first or second stimulus of the train (Fig. 1m), indicating similar short-term adaptive mechanism for the second excitatory component of the response for both conditions.

**Fig. 1.**
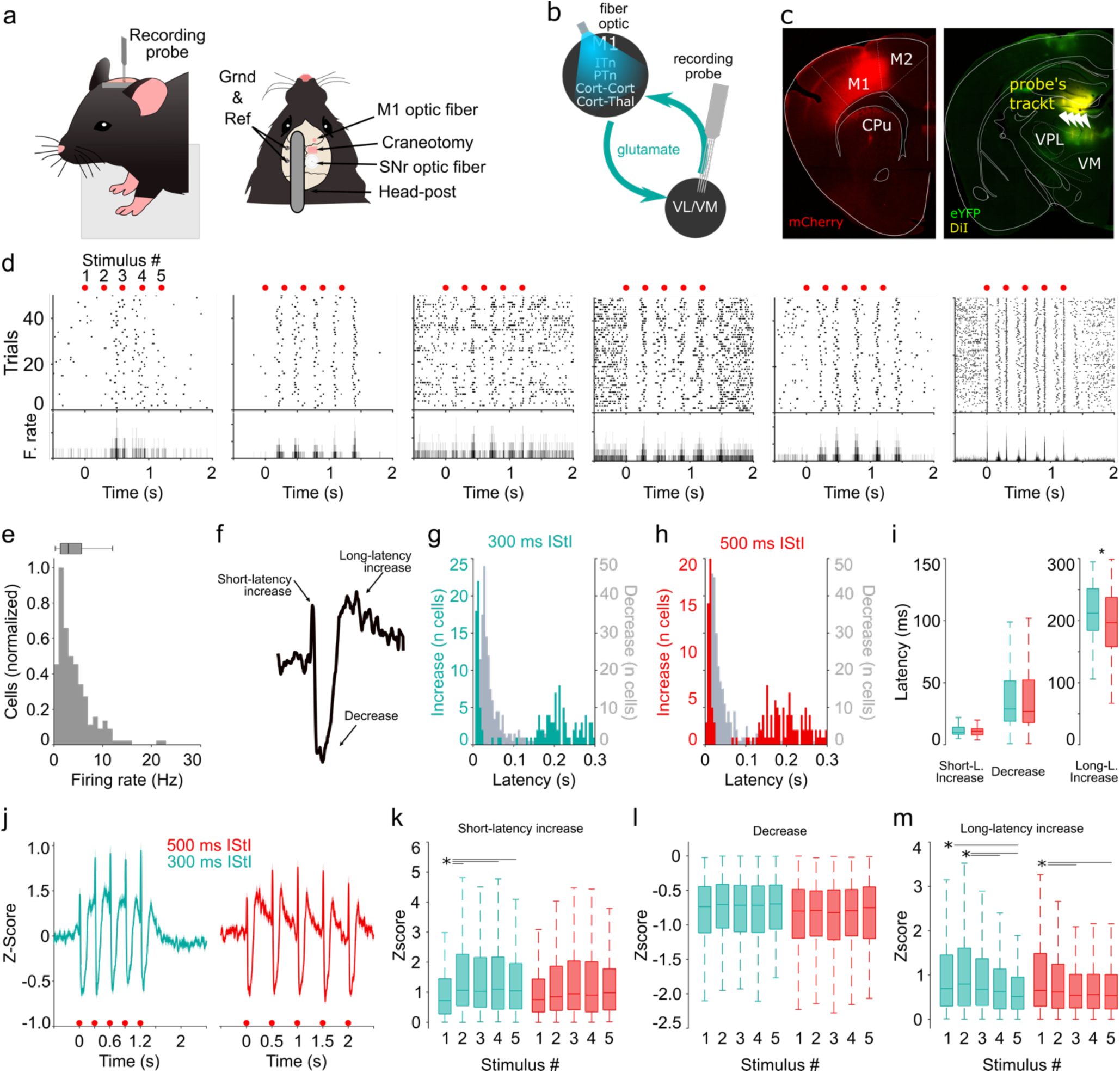
Prototypical M1-evoked responses in VL/VM. **a)** Schematic representation of the head-fixed preparation for mice. **b**) Diagram of the experimental setup in the corticothalamic circuit. M1-evoked neural activity was recorded in VL/VM. Intratelencephalic neurons (ITn), pyramidal tract neurons (PTn), cortico-cortical neurons (Cort-Cort), and cortico-thalamic neurons (Cort-Thal). **c)** Representative histological confirmation of the stimulation sites in M1 (mCherry and eYFP proteins were used as reporters) and the recording site in VL/VM. The four shanks of the silicon probe (DiI stained) are indicated with arrowheads. **d)** Representative spike rasters and their corresponding average peri-event histograms for six neurons recorded in VL/VM. Activity was aligned to the first stimulus of the train (indicated by red dots at the top). The five stimuli of the stimulation train (3.3 Hz) are indicated on top of each plot by red dots. **e)** Firing rate histogram for all cells recorded. **f)** Representative prototypical average response for the first stimulus of the train. The different components of the response are indicated. Response latency histograms for the short-latency excitatory and inhibitory components of the stereotypical response for the 300 ms (**g**) and 500 ms (**h**) inter-stimulus interval. **i**) Comparison of short- and long-latency responses between the 300 ms and 500 ms conditions. **j**) Full train-averaged population response for the 300 ms and 500 ms conditions. Comparison of the response amplitude for the short-latency (**k**), decrease (**l**), and long-latency (**m**) components of the train-evoked responses. Boxplots indicate median and 75^th^ and 25^th^ percentiles. Statistical differences are indicated by asterisks and lines joining specific comparisons (Bonferroni post hoc test, * *P < 0.05*). Mann-Whitney test was applied in I. Kruskal-Wallis [K-W] was applied in K (300 ms, degrees of freedom *[df] = 4; X*^2^ *= 14.89; p < 0.005; 500ms df = 4; X*^2^ *= 5.86; p = 0.2*), L (*df = 4; X*^2^ *= 3.42; p = 0.49; 500ms df = 4; X*^2^ *= 1.31; p = 0.859*) and M (300 ms, *df = 4; X*^2^ *= 24.86; p < 0.001; 500ms df = 4; X*^2^ *= 12.6; p = 0.013*).

Then we explored if as a population, VL/VM neurons may dynamically adapt to two different ISIs. First, we plotted the averaged (z-scored) activity of each of the 377 thalamic neurons that were active in both stimulation protocols aligned to the first cortical stimulus of the train. Neurons in both conditions were sorted according to the time of their highest firing rate between the first and second stimulus of the train in the 300 ms ISI protocol (300 and 500 ms; Fig. 2a). This representation suggested that, regardless of the ISI, the population activity organized into similar sequential activations that spanned around 300 ms after stimulus onset, even for the 500 ms ISI. This was confirmed when we plotted all neurons active on each protocol (i.e. not only those firing in both protocols) and sorted the activity of both conditions according to their own response latencies to the 300 ms or 500 ms ISI protocol (Supplementary Fig. 1a). The same kind of sequential responses were observed in a subset of experiments (3 animal, 157 neurons) where the same stimulation protocols were given at different light intensities ranging from ∼ 0.6 to 20 mW (Supplementary Fig. 2, see methods). Then, upon aligning the average population response to the onset of each stimulus of the train, we identified virtually identical responses for both ISIs (Fig. 2b, bottom left). Next, the evoked population averages were compared with population responses constructed with spike trains randomly circularized from the activity of the same neurons (see methods; Fig. 2b, bottom left, gray traces). Finally, for each neuron we calculated the correlation coefficient between the activity evoked by the 300 and 500 ms ISI trains and between the surrogated spiking activity. The correlation distributions between the 300 and 500 ms ISI trains were significantly higher than those comparing the surrogated spike trains (Fig. 2b, bottom right, gray bars). These data suggest that, as a population, the VL/VM network may not be temporally adapting to the two stimulating conditions. To further characterize this possibility, we applied a geometric approach^29,30^ to compare the matrices from our two ISI conditions. For the first comparison, called “relative” comparison, 30 bin matrices were constructed for both ISIs, that is, the bin sizes were adjusted so that both ISIs fitted 30 bins (10 / 16.6 ms bins for 300 / 500 ISIs; Supplementary Fig. 1b). For the second comparison, called “absolute” comparison, 30 bin matrices were constructed with the first 300 ms (30 bins) after the stimulus onset for both ISIs (Supplementary Fig. 1c). We averaged the activity of the five stimuli of the train, resulting in an averaged neural sequence for each ISI (Supplementary Fig. 1b-c). Then we calculated Euclidean distance matrices for each ISI and across ISIs for relative and absolute comparisons (Fig. 2c) and reported the average Euclidean distance (Fig. 2d) and diagonal asymmetry index (see methods; Figure 2e) between the absolute and relative matrices depicted in Supplementary Fig. 1b-c. Relative comparisons presented significantly higher Euclidean distances and a diagonal asymmetry index, indicating that when comparing bin by bin in real time (absolute comparison), both 300 ms and 500 ms ISIs produced indistinguishable population dynamics. Our results indicate that under these passive stimulation conditions, VL/VM appears to produce non-temporally-adapting stereotypical population responses evoked by each cortical stimulus of the train, suggesting a robust preconfigured organization in the cortico-thalamic communication.

**Fig. 2.**
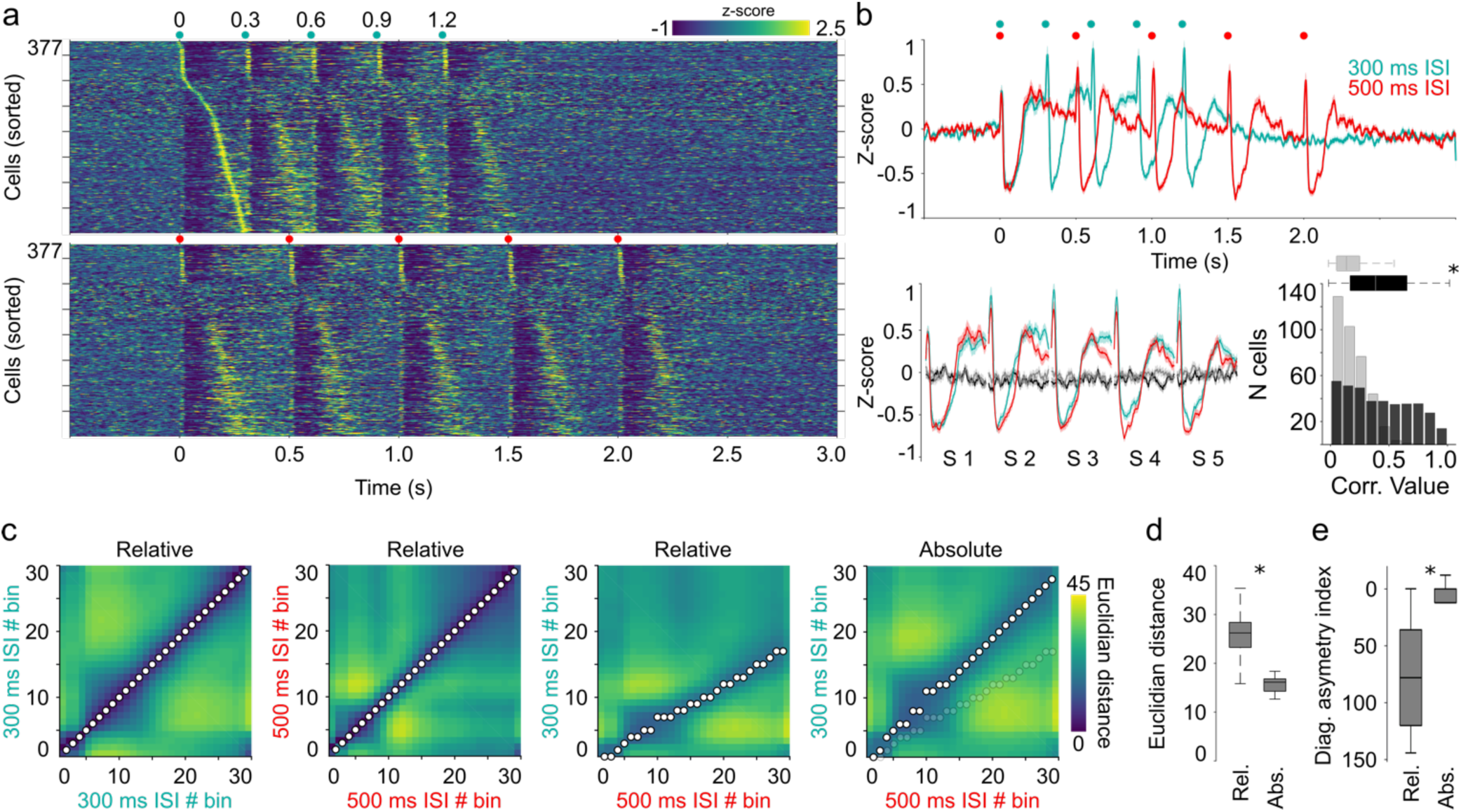
Stereotypical M1-evoked responses in VL/VM. **a)** Average firing rates (z-scored) evoked by M1 stimulation for cells recorded under two conditions: 300 ms (top) and 500 ms (bottom) inter-stimulus intervals (ISIs). Cells were sorted according to the moment of their highest firing rate between the first and second stimulus of the train. Each stimulus of the train is indicated on top of each panel (red dots). **b**) Full train-averaged population response for the 300 ms and 500 ms ISIs (top). Averaged population activity aligned to each stimulus of the train (in serial order; bottom). **c**) Euclidean distance matrices comparing 300 ms and 500 ms dynamics in the relative (time normalized) or absolute sorted matrices displayed in Figure S1B and Figure S1C, respectively. The minimum distance per bin is depicted with circles. For visual comparison, for the 300/500 ms absolute comparison, minimum distances for the relative comparison are also displayed in shaded circles. Box plot (25^th^ & 75^th^ percentiles) for the Euclidean distances (**d**) and diagonal asymmetry index (**e**) between the relative and absolute 300/500 ms comparisons. Significant differences are indicated by asterisks (* *p < 0.05*) and obtained by applying the Mann-Whitney test.

Next, we explored if the sequential stereotypical population responses could be composed of subgroups of neurons expressing specific activity patterns. To this end, we applied a principal component analysis (PCA)/Silhouette-based method of classification^31^ (see methods) on the perievent histograms obtained from the individual raster plots (Fig. 1d) over the entire population of cells and for both conditions (300 and 500 ms ISIs). We applied this method to the 377 neurons with stable responses in both protocols. Because the 300 ms ISI protocol and the first 300 ms of the 500 ms ISIs protocol produced virtually identical average responses on each stimulus of the train (Fig. 2 & Supplementary Fig. 1), we focus our analysis on the first 300 ms after each stimulus for both 300 and 500 ms data. The projection with the highest Silhouette’s values indicated that cells could be better grouped into five different types of activation patterns (pattern clusters) evoked by M1 stimulation (Fig. 3a-b) and all patterns were present in both ISIs. The first pattern (Type 1) presented very low or non-response amplitudes. This pattern captured about 45-50% of the entire population of neurons (Fig. 3c, top). The remaining four patterns consisted of short-latency transient activations followed by inhibitions, sharp activations, or inactivation followed by rebounds (Fig. 3a-b). The structure of these patterns was maintained in both ISIs and visible in the average response pattern (Fig. 3a) or in individual representative neurons (Fig. 3b), and these patterns were almost homogenously distributed in the remaining ∼50% of the population of neurons (Fig. 3c top). Then we estimated if all response patterns were present in the 18 animals recorded under these conditions. We found that pattern type 1 was present in the 18 animals, but, interestingly, pattern types 2 to 5 were also present in most of the animals (Fig. 3c, bottom). More importantly, we found that 12 out of the 18 animals presented the five patterns, two out of 18 presented four patterns, and only four animals presented only one pattern. Altogether, these results indicate that passive stimulation of M1 in awake, naïve behaving mice produces robust stereotypical patterns of thalamic activation preserved across animals, further supporting a preconfigured organization. If this were the case, it could also be expected that this organization would be present under different brain-state conditions, such as anesthesia. To explore this possibility, we used three animals that were recorded in the previously described conditions for two sessions (the animals used for light intensity experiments depicted in Supplementary Fig. 2) and in a third and last session, animals were administered with a 1 g/kg dose of the anesthetic urethane, know to preserve different sensory evoked dynamics in rats^26,31,32^. We observed that in this subgroup of animals, M1-stimulation evoked almost indistinguishable averaged population dynamics (Supplementary Fig. 3a-b) and response patterns (Supplementary Fig. 3c-e) in awake than anesthetized conditions. Here is important to highlight that the anesthetized recordings were performed after two sessions where at least 800 stimulation protocols were provided, and still, the response patterns remained stable, strongly supporting the conclusion of preconfigured response, but opening the question of its contribution during movement execution.

**Fig. 3.**
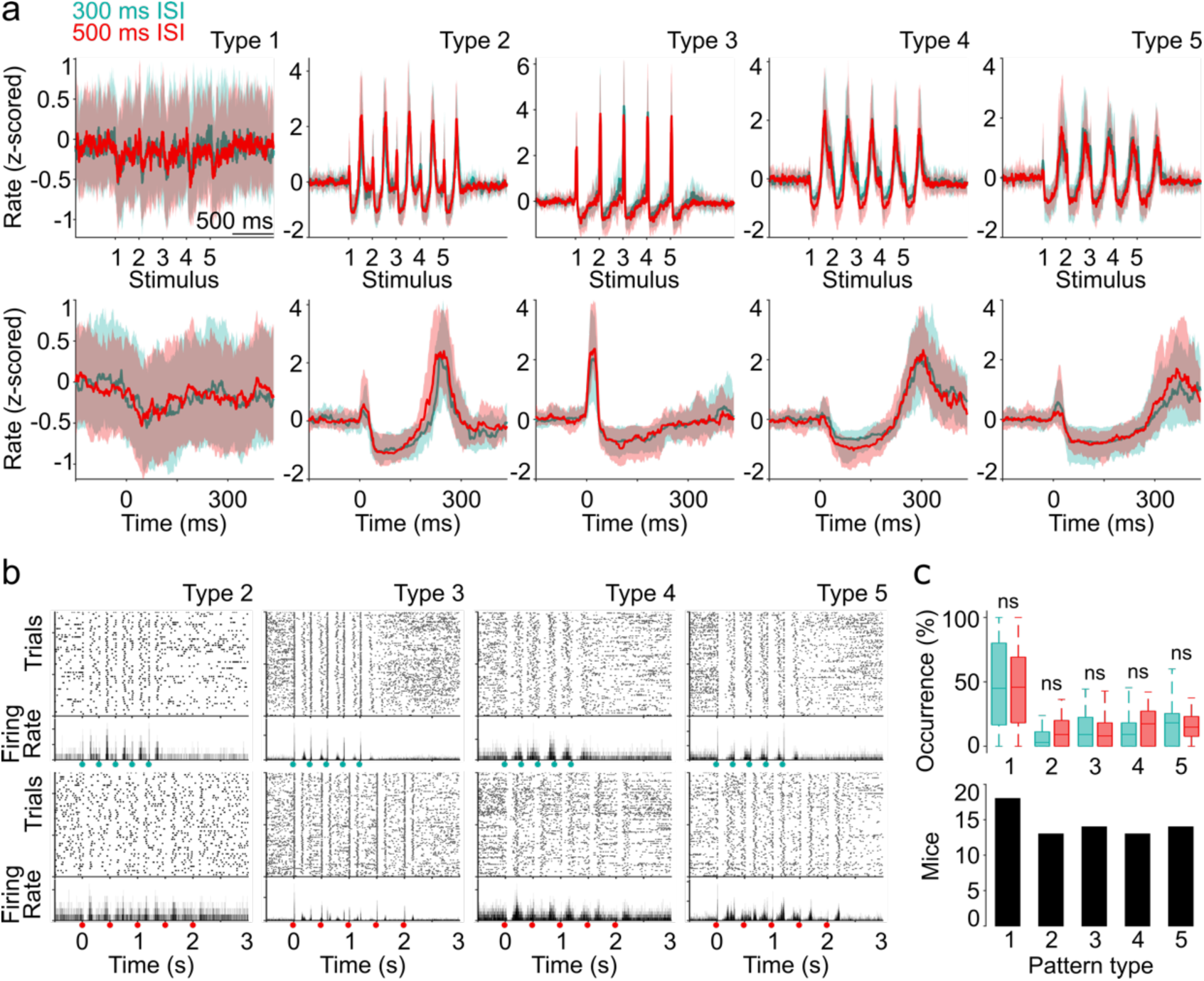
M1-evoked response patterns in VL/VM. **a)** Average M1-evoked patterns for cells classified as part of specific pattern clusters for the 300 ms (green) and 500 ms (red) conditions. The top row displays the pattern evoked by the full stimulation train and the bottom row depicts a magnification for the first stimulus of the train. Solid lines and shaded areas represent the median and the 25^th^ and 75^th^ percentiles, respectively. **b**) Representative spike rasters and their corresponding average peri-event histograms for four neurons (same neuron on each column) belonging to specific pattern clusters recorded in both the 300 ms (top row) and 500 ms (bottom row) conditions. Activity was aligned to the first stimulus of the train (indicated by green and red dots for 300 and 500 ms ISI, respectively). **c**) Percentage of cells belonging to each pattern cluster displayed in A (upper panel). Number of animals that presented each pattern (lower panel). Boxplots indicate median and 75^th^ and 25^th^ percentiles. Statistical comparisons were performed by applying K-W and Bonferroni post hoc tests. In panel C, *df = 9; X*^2^ *= 65.66; p < 0.001*.

### Motor thalamus responses associated to movement execution

Next, we explored whether the M1-evoked thalamic nPAPs described in the previous sections could be related to specific parameters of movement execution. With this objective, we established a behavioral protocol compatible with our head-fixed recording setup. Mice (n = 13) were trained to handle a lever located under their left forepaw, contralateral to the recording sites (Fig. 4a). Behavioral protocols were divided into three phases (Fig. 4a, right). In the first phase, named “Handling and Modeling”, the animals were accustomed to head-fixing conditions for about 2-3 sessions (20-30 min per session, 1 session per day). Immediately after, animals were water-restricted and started a modeling period where they learned to displace the lever to obtain rewards (water drops delivered through the waterspout). The modeling period lasted about 10 sessions (50-60 min sessions, 1 session per day), where a reward was delivered with progressively longer movements, starting from 50 ms displacements (0.1 mm displacement threshold) to 500 ms displacements (0.8 mm displacement threshold). We rewarded movements in any possible direction, but individual animals quickly displayed a preference for either backward or forward directions (i.e., pushing or pulling displacements, respectively). Lever trajectories were constantly recorded, analyzed, and aligned to reward delivery (Fig. 4a), indicated by a 1 s green light located in the front of the mice. After the modeling period, animals were changed to the formal training in the two-duration version of the task. Here mice were asked to perform 200 or 500 ms movements in alternating blocks of 20 consecutive trials for each duration. The 200 and 500 ms blocks were signaled by a contextual white or blue light, respectively. Movement duration was selected for two reasons: to match as closely as possible the two ISIs used in the passive stimulation experiments and to analyze two movements with at least twice the difference between them. Learning curves were constructed for different movement parameters. First, we calculated the intralimb variability by computing the Pearson correlation coefficient between each possible pair of trajectories for each category (200, 500 ms; Fig. 4c) as well as its variance (Fig. 4d). These parameters are indicators of how stable the movement trajectory is over the course of learning. For both parameters we observed a quick improvement in the modeling sessions that remained stable during the rest of the training. Consistent with an increase in difficulty, we also observed that the 500 ms movements produced significantly lower correlation values and higher variances, especially in the early sessions of the two-duration phase (Fig. 4b-d). Then, we calculated the movement overshoot, defined as the amount of time that the animals maintained the movement after reward delivery. This measure indicates movement duration. We observed that animals presented similar overshoot values for the two durations, rounding about 200 ms (Fig. 4e). An indication of movement efficiency is the number of attempts that the animals make to obtain a single reward. To estimate this value, for each trial we quantified the cumulative amount of time that the lever was pressed to obtain the reward. Consistent with motor improvement, this value decreased significantly with training, even for the more difficult 500 ms movement (Fig. 4f). Movement speed maintained similar values throughout the learning curve and different protocols (Fig. 4g).

**Fig. 4.**
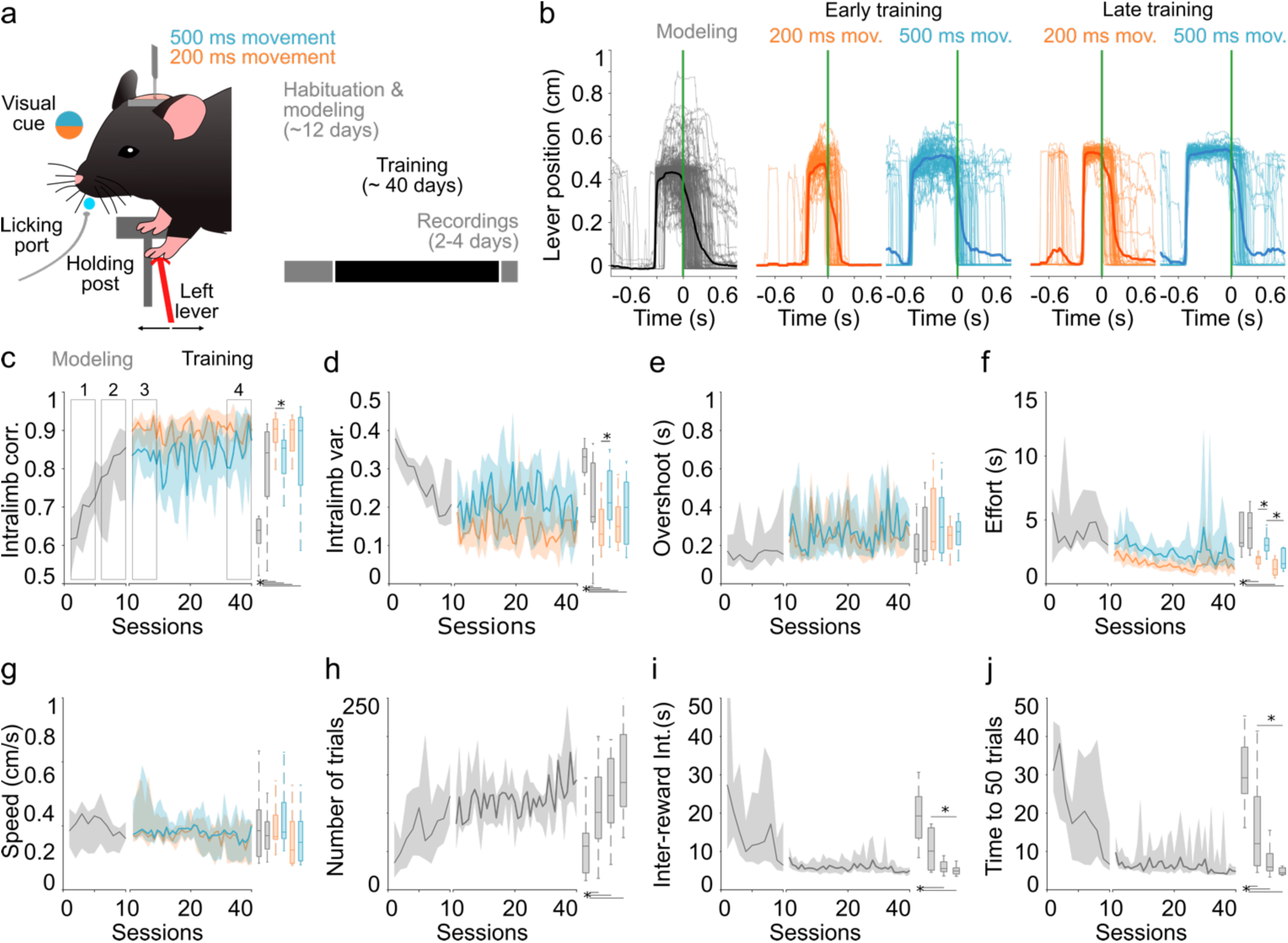
Behavioral protocol. **a)** Schematic representation of the behavioral setup. **b**) Representative trajectories for trials in the 200 ms and 500 ms blocks in difference stages of learning. (**c** to **j**) Learning curves (left panels) and boxplot comparisons for specific groups and sessions (right panels) including the modeling phase (gray) and the training phase in the two-interval version of the task (color coded) for the following variables: Intralimb correlation **(c**), variability (**d**), overshoot (**e**), effort (**f**), speed (**g**), number of trials (**h**), inter-reward interval (**i**), and time to reach the first 50 trials (**j**). Data is presented as median (solid line) ± 75^th^ and 25^th^ percentiles (shaded area). Boxplots indicate median and 75^th^ and 25^th^ percentiles for groups of five sessions at the four stages of the learning curves, indicated by the numbered rectangles in C. Statistical differences are indicated by asterisks and lines joining specific comparisons (Bonferroni post hoc test, P < 0.05). K-W values for C (*df = 5, X*^2^ *= 31.76, p < 0.001*), D (*df = 5, X*^2^ *= 25.9, p < 0.001*), E (*df = 5, X*^2^ *= 5.15, p = 0.398*), F (*df = 5, X*^2^ *= 30.48, p < 0.001*), G (*df = 5, X*^2^ *= 4.43, p < 0.488*), H (*df = 3, X*^2^ *= 16.41, p < 0.005*), I (*df = 3, X*^2^ *= 21.73, p < 0.001*), J (*df = 3, X*^2^ *= 26.87, p < 0.001*)

Finally, we estimated three more general performance values, the total number of trials performed in each session (Fig. 4h), the inter-reward interval (Fig. 4i), and the total amount of time to reach the first 50 trials, which is the period where animals are more engaged in the task. (Fig. 4j). All those values significantly improved with training. Altogether, the previous results demonstrate a robust behavioral protocol where neural data can be associated with two different movements matching our stimulation protocol in the temporal domain. Once animals were well-trained in the task, a craniotomy to access the VL/VM was performed (see methods). The following day, the animals resumed training for 3-4 days and were recorded (same as in the previous sections; one recording per session/day) while performing the behavioral protocol.

Under these conditions we were able to record 461 neurons from 13 mice during 36 sessions (2 to 4 recording sessions per animal). A great variability of activity patterns was observed when aligning the neural activity to the moment of reward delivery (Fig. 5a). These patterns were visible in both the 200 and 500 ms movement trials. Then, we explored if the spiking activity of individual neurons could be linearly associated with different parameters of movement execution, such as speed, overshoot, or amplitude, but found no evidence of such relationship (Supplementary Fig. 4; see methods). As a population, the activity of the cells was organized as a gradient of activation/inactivation with at least two visible ends. On one end, a group of neurons decreased their firing activity during movement, and on the other end, a group of neurons visibly increased their spiking activity during movement (Fig. 5b). Interestingly, this structure remained almost unchanged in the 200 and 500 ms movement trials (Fig. 5b; both matrices were sorted from highest to lowest average firing rates in the 200 ms condition). When averaging neuronal activity in both types of trials (Fig. 5c), we divided the population dynamics into three phases: a rising phase that reached its highest peak before movement onset in either type of trial (indicated by color-coded arrows in Fig. 5c); a descending phase that was interrupted by the reward delivery; and then, a transitory, sharp response that was almost identical in both types of trials. To further explore potential differences between the neural activity in the 200 and 500 ms trials, for each neuron, we calculated the Pearson correlation coefficient between the average response during the 200 and 500 ms trials. In this case, we normalized the temporal domain to have a similar number of bins in both conditions. Hence, we observed that most of the cells displayed high correlation values, indicating the maintenance of the spiking pattern independently of the movement duration (Fig. 5d). The distribution of correlation values was also compared with a surrogated distribution obtained by randomly circularizing the spiking activity of each neuron (see methods). Because average calculations over the whole population may mask the diversity of activation patterns (Fig. 5a), we used the same PCA/Silhouette-based method to detect specific optogenetic-evoked patterns in our previous sections (Fig. 3), but in this case to identify potential behaviorally related patterns. As before, we normalized the temporal domain to compare activity in both types of trials. Our analysis indicated that, based on the Silhouette’s values (Supplementary Fig. 5a), neurons could be grouped into 10 patterns (Fig. 5e), where pattern type 1 was not particularly associated with an increase or decrease in activity; hence, we classified these neurons as non-related to the task. The remaining patterns confirmed the presence of widely diverse activity patterns surrounding movement onset and reward delivery. With subtle changes, these patterns reflected subregions of the general architecture of the average population response depicted in Fig. 5c. Some patterns presented ramping activity with the highest peaks around movement onset (Fig. 5e, types 4 and 6 and 7); other groups presented different levels of transient inactivation (Fig. 5e, types 2, 5, 8 and 9); and other groups of neurons presented increased activity triggered by reward onset (Fig. 5e, types 3 and 10). Interestingly, while the highest Silhouette values were found for the 10-cluster projection, a four-cluster projection also presented significantly high values (Supplementary Fig. 5a-c), and these projections also matched the general architecture of the average population response. In terms of proportions, the non-responsive neurons comprised the biggest group, and the rest of the patterns were integrated by a similar number of neurons (Fig. 5f). To estimate if a particular pattern was preferentially associated with the 200 or 500 ms movement trials, we selected sessions where at least 8 neurons were recorded simultaneously (24 out of the 36 sessions) and estimated the prevalence of each pattern in each session. We found that all patterns presented a similar prevalence during the 200 and 500 ms trials (Fig. 5g). The fact that the two types of trials produced indistinguishable VL/VM population dynamics suggests that the duration for this type of movement is not particularly encoded in this region. On the other hand, this data also suggests that while different patterns of activation could be formally extracted, it appears that thalamic neurons adjusted to three main phases: an ascending phase, roughly finishing around movement onset, followed by a descending phase, and an abrupt increase triggered by reward.

**Fig. 5.**
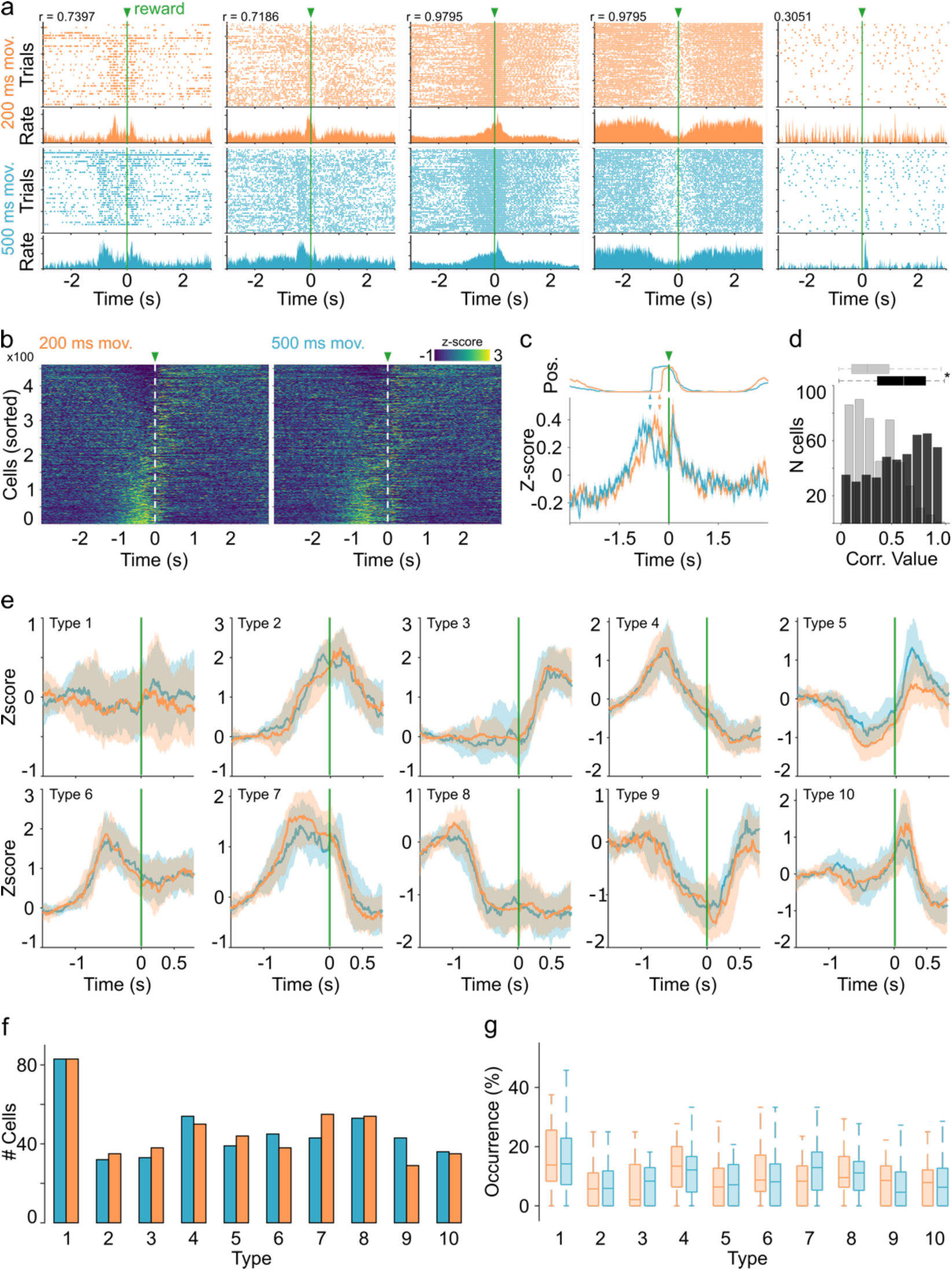
Thalamic signals during movement execution. **a)** Pairs of spike rasters (top) and average perievent histograms (bottom) for five representative cells with different response patterns recorded during the execution in the two-interval task during the 200 ms (top row, orange code) and 500 ms (bottom row, blue code) trials. Spiking activity was aligned to the reward onset (green lines and arrowheads). **b)** Z-scored averaged activation of the firing rate of all recorded cells during the 200 ms (left) and 500 ms trials. Cells are sorted according to the average maximum firing rate between -1.2 s and reward delivery, in zero, and indicated by a green arrow and with a dotted line. **c)** Averaged perievent histogram of the firing rates (bottom) for all cells recorded during the 200 ms (orange) and 500 ms (blue) trials aligned to the reward delivery (green line and arrow). The averaged lever trajectories for both types of trials is presented on top of the plot. Color-coded arrows indicate movement onsets. **d)** Pearson correlation coefficients between normalized averaged patterns from 200 and 500 ms trials for all recorded neurons (black). Surrogate data is presented in gray. Boxplot indicates median and 75^th^ and 25^th^ percentiles, *\* Mann-Whitney test, p < 0.05*. **e**) Average behaviorally evoked perievent histograms for cells classified as part of specific patterns for the 200 ms (orange) and 500 ms (blue) conditions. **f)** Absolute number of cells expressing each pattern. **g)** Percentage of cells belonging to each pattern. Percentages were calculated using sessions with more than 8 simultaneously recorded cells. K-W values (*df = 19, X*^2^ *= 41.29, p = 0.002*).

### M1-evoked thalamic nPAPs are linked to specific movement-related activity patterns

Next, we ask whether the activity of the neurons related to different phases of the movement could also be related to the preconfigured patterns of activation triggered by M1 stimulation. To this end, immediately after each behavioral session, we performed the same M1 stimulations as the the ones performed in our naïve animals (Fig. 1-3). In this way, we were able to identify behaviorally related and M1 stimulation-related activity in the same neurons. Because our previous analyses showed no differences between the 300 and 500 ms ISIs (Fig. 2-3), for the following analysis, we focused on the 300 ms ISI train. As in the naïve animals, VL/VM neurons recorded from expert animals also displayed robust activation patterns in response to M1 stimulation (Fig. 6a, left panel). Then, to compare these patterns with those originally reported for the naïve animals, we pooled all neurons recorded from both groups (naïve and expert) and re-applied our PCA/Silhouette-based method to identify groups of neurons with similar response patterns. Here again we found the five patterns observed in Figure 3. Both groups of animals presented neurons in the five groups, further confirming the strong preconfigured organization of this cortico-thalamic interaction (Fig. 6b). However, we also observed that pattern type 2 was significantly more represented in the expert group (Fig. 6b, bottom right panel). Once it was confirmed that the neurons in the expert animals also displayed archetypical responses to M1, we explored potential crossings between behavioral and M1-related representations. In our first approach, we sorted the activity of the cells evoked by M1 stimulation but according to the order obtained from their activity during the behavioral sessions for the 200 ms movement trials displayed in Figure 5b. This representation showed that cells that increased their firing rate during behavioral sessions appeared to produce more robust responses to M1 stimulation than the cells that decreased their firing rate during behavioral sessions (Fig. 6a, right panel). To confirm this possibility, we selected the neurons located at both ends of the distribution produced during behavioral sessions (Fig. 5b), that is, cells below and above the 25^th^ and 75^th^ percentiles of the distribution, respectively. The average subpopulation response of these groups confirmed their characteristic activation and inhibition profiles during movement execution (Fig. 6c). But most importantly, when plotting their activity evoked activity during M1 stimulation, responses were clearly different between groups (Fig. 6d). Neurons with increased responses during behavior displayed higher amplitudes for the excitatory and inhibitory components of the M1-evoked response. Then, we analyzed if these two groups would also differ in terms of short-term adaptation to the stimulation train. To this end, we focused on the increase in long-latency firing rate (between 100 and 290 ms after each stimulus). We found that the group associated with increased responses during behavior presented a strong adaptation as the stimuli progressed, while the group associated with inhibitions presented little evidence of adaptation (Fig. 6e). This could be related to the fact that the responses of this group in the extreme of the distribution are mainly composed by pauses in activity. To confirm that these differences were related to the lower and upper ends of the behavioral distribution, we performed the same analysis but in cells from percentiles 35 to 50 and 50 to 65, where behaviorally related activity was more homogenous (Fig. 6a, right panel). In this case, both groups of neurons presented similar patterns of activation and adaptation (Fig. 6f-g). Finally, we explored the possibility of a more specific relationship between behaviorally related and M1-evoked responses. For this, we tried to link PCA/Silhouette-based patterns obtained during behavior (the 10 different pattern clusters from Fig. 5e) with those obtained by M1 stimulation (Fig. 6a; five different pattern clusters). To perform statistical comparisons, we selected sessions with at least 10 simultaneously recorded neurons, and then we calculated the prevalence of M1 stimulation patterns for each of the behaviorally related patterns. While this high number of possible combinations decreased the statistical power, we were still able to detect that for behaviorally related pattern #2 there was a significantly higher prevalence for M1-related pattern #1 (Supplementary Fig. 5d). On the other hand, behaviorally related pattern #5 also displayed a significant bias for M1-related pattern #4 (Supplementary Fig. 5d). These results demonstrate that M1-evoked preconfigured responses are directly linked to thalamic responses associated with specific phases of forelimb movements.

**Fig. 6.**
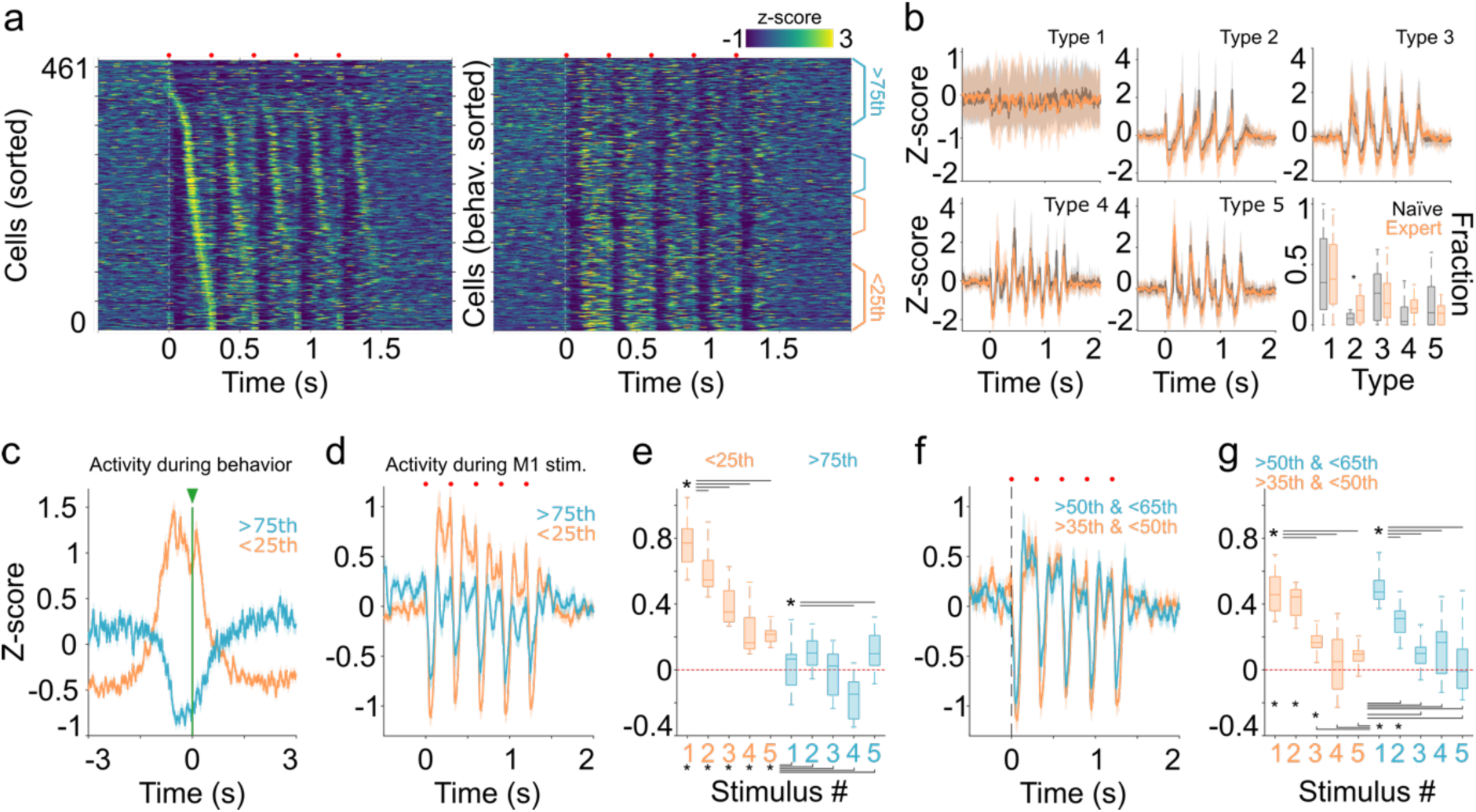
Stereotypical thalamic patterns constrain movement related activity. **a)** Average firing rates (z-scored) evoked by M1 stimulation for all cells recorded from experienced animals. Cells were sorted according to their highest firing rate between the first and second stimulus of the train (left) or according to the sorting order obtained from their activity during behavioral execution displayed in Figure 5B (right). **b**) Average M1-evoked patterns for cells classified as part of specific pattern clusters for the 300 ms in trained animals (orange); these patterns were compared with those obtained from naïve animals (gray). Solid lines and shaded areas represent the median and the 25^th^ and 75^th^ percentiles, respectively. Bottom right panel display the percentage of cells belonging to each pattern for each group, *K-W values* (*df = 9, X*^2^ *= 58.45, p < 0.001*). **c**) Average firing rate of two subgroups of cells during behavioral execution and belonging to above and below the 75^th^ and 25^th^ percentiles of the sorted distribution, respectively. The sorting order was obtained during the behavioral execution displayed in Figure 5B (also indicated in orange and blue in the right matrix of panel A). Activity is aligned to reward delivery in zero (green line and arrow). **d**) Average firing rate evoked by M1 train stimulation (indicated by red dots) for the same subpopulations. Activity is aligned to the onset of the first stimulus of the train in zero. **e**) Amplitude of the subpopulation late responses for each stimulus of the train, *K-W values* (*df = 9, X*^2^ *= 1167.69, p < 0.001*). **f** & **g**) Same as in D and E but for subpopulations belonging to the 50^th^ to 65^th^ and 35^th^ to 50^th^ percentiles of the population (as indicated in A, right) *K-W values* (*df = 9, X*^2^ *= 968.11, p < 0.001*). Boxplots indicate median and 75^th^ and 25^th^ percentiles. Statistical differences are indicated by asterisks and lines joining specific comparisons (Bonferroni post hoc test, *P < 0.05*). Red dotted line in E & G is depicted as visual reference.

The previously described phenomenon opens two questions. First, under which circumstances would these preconfigured responses be changed, modulated, or abolished? And second, would nPAP modifications impact behavior? To start answering these questions, we studied the potential modulation of the BG output, the SNr, which is one of the main inputs to the VL/VM network, over the stereotypical M1-VL/VM responses (Fig. 7a). Previous anatomical and functional observations indicate that the SNr GABAergic projections modulate the activity of VL/VM with behavioral consequences^19^. Hence, we performed the same recording configuration (M1-VL/VM) but paired it with optogenetic manipulations of SNr GABAergic neurons (Fig. 7a). In principle, this manipulation could disrupt M1-evoked thalamic nPAPs and potentially associated behaviors. To explore this possibility, in a group of Vgat-Cre animals, we expressed ChR2 (n = 4) to stimulate SNr-projecting neurons. We found that SNr/M1 paired stimulation induced a disorganization of the VL/VM responses visible at the individual (Fig. 7b) and population levels (Fig. 7c). The stimulation diminished the sharp short-latency responses but, interestingly, maintained the inhibitory component of the M1-evoked responses (Fig. 7c-d). This observation suggests that the stereotypical response is composed of an M1 sharp component, sensible to SNr modulation, and inhibitory and rebound effects, both insensible to SNr modulation. This was confirmed when we analyzed the amplitudes of the short and long-latency components of the responses. Here, SNr stimulation induced a significant reduction of the sort-latency component (Fig. 7e left) of the response, especially the 3^rd^ to last stimuli of the train, but it produced no changes in the amplitudes for the long-latency responses (Fig. 7e right). To explore potential differences in population dynamics, we constructed 30 bin matrices with the averaged activity of the five stimuli of the train, resulting in an averaged neural sequence for each experimental condition (Supplementary Fig. 6). Then, we compared the Euclidean distances between population matrices obtained from 300 vs 500 ms ISIs and the 300 ms condition with paired or unpaired SNr stimulation (Supplementary Fig. 6). The highest Euclidean distances were observed for the latter comparison, confirming that SNr modulation significantly modified the VL/VM population representation of the M1 stimulus (Fig. 7f). Next, we explored if the average/population changes were related to a particular subgroup of neurons. To this end, we used our PCA/Silhouette-based classification method and selected the best projection with five groups. This projection yielded five almost identical response patterns to the ones obtained in the original analysis of Figure 3 (Fig. 7g). In this case, SNr stimulation significantly decreased the occurrence of pattern type 2 and increased the occurrence of pattern type 4 (Fig. 7h). The former was characterized by neurons with sharp, short-latency responses, while the latter was characterized by strong pauses in activity. It is important to notice that, despite the occurrence differences, all patterns were present in the four animals recorded under these conditions (Fig. 7i). The previous results indicate that the preconfigured M1-VL/VM dynamics may be partially altered by the activation of the output nuclei of the BG. Hence, to evaluate the behavioral impact of disturbing VL/VM activity, we performed SNr stimulation (directly on the SNr n =2; or its terminals in VL/VM n=2; see methods) during movement execution in highly trained animals in our two-interval task (Fig. 7j; n=4). We found that closed-loop activation triggered by minimum displacement of the lever induced significant decreases in movement stability (intralimb correlation) while sparing the rest of the variables (Fig. 7k). These effects were similar for the 200 and 500 ms movements. Altogether, the previous section suggests that altering the preconfigured dynamics in VL/VM is sufficient to produce specific deficits in movement control, in this case, movement variability.

**Fig. 7.**
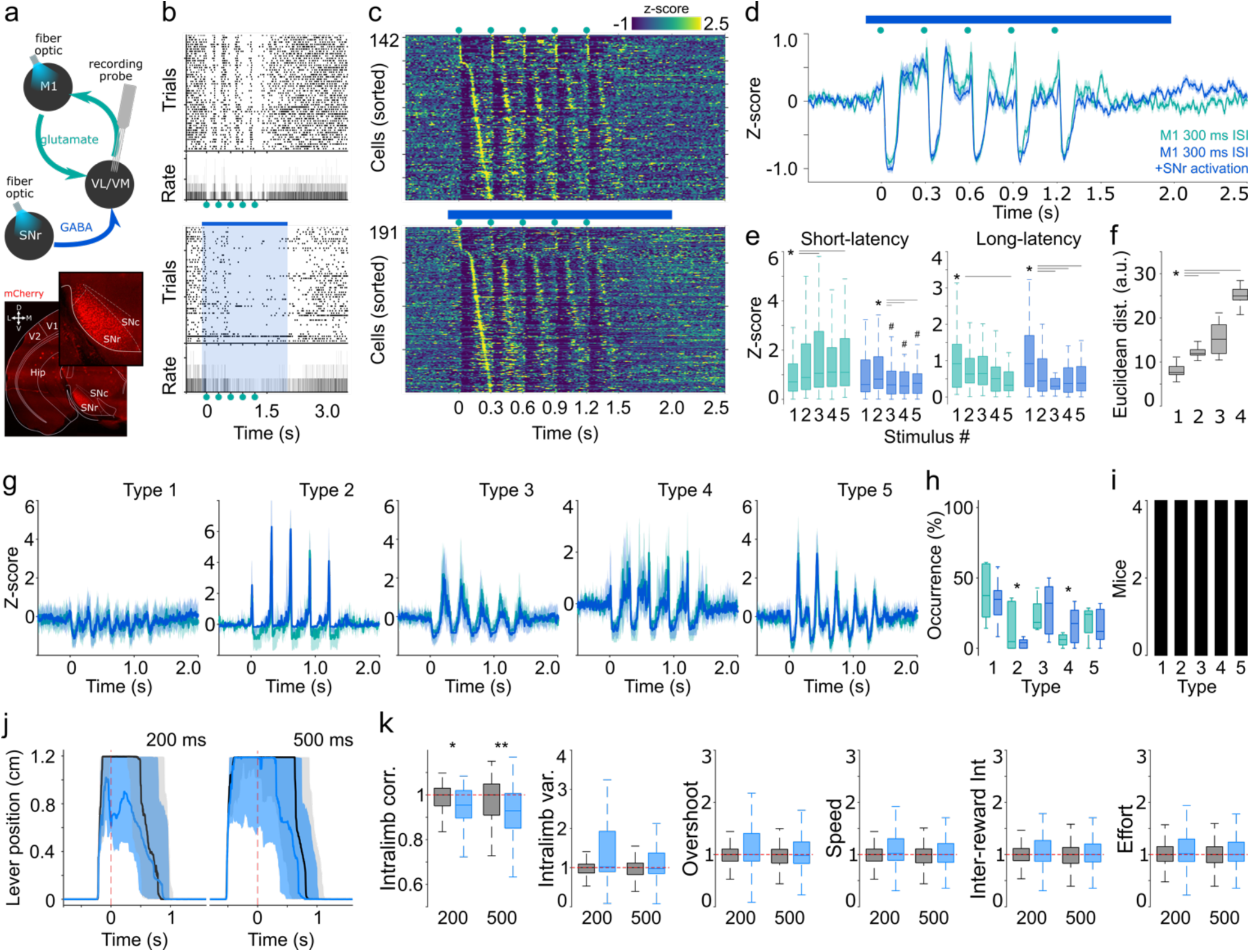
Nigral modulation of thalamic preconfigured patterns and forelimb movements. The M1-evoked responses (300 ms ISI) are compared in two conditions, with (blue code throughout the figure) and without (green code throughout the figure) SNr paired stimulation. **a**) Experimental configuration where M1-evoked neural activity was recorded in the VL/VM region with SNr simultaneously stimulated (top). Histological confirmation of the stimulation sites in SNr. **b**) Representative spike rasters and their corresponding average peri-event histograms for a neuron recorded in the VL/VM with (bottom) and without (top) associated SNr stimulation (blue bar and shade). Activity was aligned to the first stimulus of the train (indicated by green dots). **c**) Average firing rates (z-scored) evoked by M1 stimulation for cells recorded in a group of naïve animals under two conditions: M1 stimulation (top; indicated by green dots) and M1 stimulation paired with SNr stimulation (bottom; indicated by blue bar). Cells were sorted according to the moment of their highest firing rate between the first and second stimulus of the train. **d**) Average M1-evoked average firing rates for cells recorded with (blue) and without (green) paired nigral stimulation (same color code for the rest of the figure). **e**). Comparison of the response amplitude for the short-latency (left panel; *K-W values, df = 9, X*^2^ *= 51.58, p < 0.001*), long-latency (right panel; *K-W values, df = 9, X*^2^ *= 50.57, p < 0.001*) components of the train-evoked responses with and without paired nigral stimulation. **f**) Euclidean distance comparison for M1-evoked sequential activation matrices (30 bins, 5 stimuli averaged) for 300 ms ISI trains with and without SNr paired stimulation. Comparison performed between the following conditions: 300 ms and 500 ms (absolute first 300 ms; **1**); 300 ms and 500 ms (first 300 ms) ISI with SNr paired stimulation (**2**); 300 ms and 500 ms (time normalized; **3**); and 300 ms ISI with and without SNr paired stimulation (**4**). *K-W values, df = 3, X*^2^ *= 100.95, p < 0.001.* **g**) Average M1-evoked patterns for cells classified as part of specific patterns for the 300 ms in both conditions. Solid lines and shaded areas represent the median and the 25^th^ and 75^th^ percentiles, respectively. **h**) Percentage of cells belonging to each pattern in both conditions. **i**) Number of animals that presented each pattern. *K-W values, df = 9, X*^2^ *= 28.93, p < 0.001.* **j**) Representative average trajectories (shaded areas, 25^th^ and 75^th^ percentiles) of the lever position aligned to the reward delivery (time “0”, red dashed line), for stimulated (blue) and non-stimulated (black) trials during 200 ms (left) and 500 ms (right) trials. **k**) Box plots (25^th^ and 75^th^ percentiles) displaying group comparisons (n = 4) between non-stimulated (black) and stimulated (blue) trials. For each variable (indicated in the “Y” axis), data for each animal was normalized to the median of their control condition. Significant differences are indicated by asterisks (* *p < 0.05*; ** *p < 0.01*) and obtained by applying Kruskal-Wallis and Bonferroni post hoc tests (K-W, for intralimb correlation, *df = 3; X*^2^ *= 13.26; p = 0.004; for the rest of the variables df =3; X*^2^ *< 5; p > 0.2*).

## Discussion

The existence of preconfigured neural dynamics has been previously reported for sensory cortices and cognitive regions, such as the hippocampus ^2^. However, until now these dynamics have not been explicitly explored in motor networks. Here we focused on the motor thalamus of awake behaving, head-fixed mice to address the possibility that preconfigured dynamics constrain movement and its related activity. Our results can be divided into two main sections: in the first study, where we describe for the first time the existence of VL/VM thalamic nPAPs a in response to M1 cortical stimulation. In the second part, we described how those dynamics are linked to behaviorally associated thalamic activity. First, in naïve behaving mice, briefly accustomed to our recording conditions, we found that cortical inputs to the VL/VM evoked complex stereotypical responses organized as transient activations and inactivations with a variety of response latencies (Fig. 1). At first sight, these patterns were very similar to the ones previously reported for the sensorimotor cortex, striatum, and SNr in anesthetized rats ^9,11,33^, suggesting a common responsive organization distributed throughout the thalamo–cortical-BG sensorimotor loops. These responses could be classified into four stereotypical patterns present in a large proportion of the VL/VM population and observed in virtually all awake (Fig. 3) and anesthetized (Supplementary Fig. 3) recorded animals. At the population level, these dynamics were organized as neuronal sequences spanning for a couple of hundred milliseconds after cortical stimulation onset (Fig. 2) and resembled dynamics previously observed in cortical and subcortical sensorimotor systems ^13,15,34,35^, referred to as “information packets” ^2^. However, contrary to what has been observed for sequential activations in the cortex and striatum ^6,36^, VL/VM dynamics were unable to dynamically adapt to different temporal intervals (Fig. 2). This result suggests that VL/VM responses to M1 activation are rigid and further supports their preconfigured nature. But what would be the consequence of this M1 stimulation-induced functional segregation? To answer this question, first we showed that during the behavioral execution of a simple forelimb movement (Fig. 4), thalamic activity can also be segregated into specific populations associated with distinct phases of the movement (Fig. 5). But most importantly, specific subpopulations identified by their response pattern to M1 stimulation were associated with specific patterns of movement-related activity (Fig. 6). These results confirm that preconfigured functional segregation, revealed by passive stimulation of M1, constraints movement-related activity in the motor thalamus. However, it is not clear if this motor-related thalamic activity is merely a reflection of M1 activity or, on the contrary, if it has specific effects on the final motor output. To clarify this point, we aimed to modulate preconfigured thalamic neural patterns by activating the SNr, a major inhibitory input to the VL/VM (Fig. 7). Interestingly, this manipulation was sufficient to modulate the early, short-latency, and excitatory components of the average population response and alter the occurrence of thalamic nPAPs. Furthermore, the same manipulation was also sufficient to significantly modify contralateral forelimb movement stability (Fig. 7).

An important open question is to determine the source of the segregated subpopulations described here. Our results suggest that this segregation would not be related to experience, since all stereotypical patterns were observed in most of the animals and in both groups, naïve and trained. Further supporting this conclusion, these patterns were also observed in urethane anesthetized conditions, demonstrating their brain-state independence, but is also worth mentioning that the anesthetized recordings were performed after two awake sessions where stimulation protocols were repeated hundreds of times. Even in these conditions, nPAPs remained stable. The possibility that these functionally segregated populations are experience-independent is consistent with recent studies performed in different non-motor thalamic nuclei, that have convincingly demonstrated the existence of genetically defined neural subpopulations associated with different anatomical profiles and functions ^37,38^. For example, a recent study ^39^ showed that the reticular thalamic nucleus, typically conceived as the GABAergic homogeneous nucleus, is composed of at least two genetically defined subpopulations with distinct functional properties. Similar results have been observed in the parafascicular thalamic nucleus ^40^, where at least three independent thalamo-cortico-striatal motifs could be functionally and genetically identified. The previous literature demonstrates an unexpected level of neural heterogeneity that could at least partially explain our results in the VL/VM. In this context, it is possible that the different nPAPs evoked by M1 stimulation are associated with genetically distinct subpopulations, like the ones previously reported for the motor thalamus ^41^. This possibility would further confirm the preconfigured nature of these signals and will be explored in future experiments. On the other hand, the thalamic nPAPs reported here may be triggered by two cortical subpopulations: pyramidal tract neurons — known to also project to multiple subcortical regions besides the motor thalamus, including the striatum and the spinal cord— and cortico-thalamic neurons projecting only to the thalamus ^42^. In this work we did not design experiments to explicitly address these two projections. However, future research will seek to address the exact contribution of these subpopulations to the nPAP architecture. Another important observation is that we were able to match specific M1-evoked patterns with specific activity associated with forelimb movements (Figs. 6, 7, Supplementary Fig. 5). Previous studies in sensory cortical regions have demonstrated that spontaneously organized patterns of activity (potentially preconfigured) are recruited by sensory stimulation ^43^. More specifically, in a recent study, researchers observed that hippocampal neurons born on the same developmental day shared relevant encoding features during adulthood, such as anatomical connectivity patterns of even spatial representations ^44^. Our data expands the interpretation of inside-out brain organization (Buzsáki, 2019) from the cognitive to the motor sphere, at least for simple forelimb movements.

Our observations also open another important question in the context of macro-circuit cortical, BG, and thalamic communication. Although the input nucleus of the BG (i.e., the striatum) is massively innervated by the cortex, the BG exerts indirect influence over the cortex because of the thalamus (VL/VM). Hence, how do the different VL/VM neural patterns interact with BG signals to eventually impact behavior? Here we started exploring this interaction by directly activating SNr-to-VL/VM projections. To our surprise, the effect of this manipulation was almost restricted to the first part of the thalamic responses, the short-latency response (Fig. 7), resulting in a significant reduction in the occurrence of one neural pattern. This observation and previous reports studying sensory cortico-thalamic loops^23,45^ suggest that the composition of M1-evoked thalamic nPAPs most likely involves a much more complex network of thalamic inputs, such as the reticular thalamic nucleus and/or the different cortical subpopulations projecting to the thalamus, both unexplored in this work. This manipulation was also associated to the disruption of movement variability. This could be considered a subtle behavioral effect; however, these results are in accordance with two main lines of evidence from the previous literature. First, M1 has been typically associated to the production motor plans, that is, stablishing the path, direction, and trajectories of movements^46–49^, while the motor thalamus has been proposed to gate and transmit information from subcortical circuits to maintain and re-organized cortical activity and associated motor commands^50,51^. Hence, is not strange to find that VL/VM manipulation by activation of SNr inputs, affected only the variability of movement and not any other parameter, such as speed or direction. In fact, recent advances in BG literature suggest that one of the main roles of these nuclei, including its output nuclei, is to modulate commanded movements^48,52–55^. Our results, however, suggest that greater efforts will be needed to fully understand how the complex activity patterns observed at the single and population levels in the striatum will be translated into the VL/VM and eventually to the motor cortex. For example, it is still difficult to picture how start, stop, speed, and time signals that are consistently recorded throughout the BG ^6,9,56^**^‒^**^60^ would be transformed or simply relayed by the VL/VM to the motor cortex. Our results suggest that upcoming signals from subcortical regions, such as the SNr, will interact with rigid, stereotypical patterns of thalamic activity. How these interactions will be translated into organized motor commands or modulation of specific movement parameters will be the subject of future investigations. On the other hand, while we found that trained animals presented discrete differences with respect to naïve animals, the effects of learning on cortico-BG-thalamic dynamics must be thoroughly and explicitly explored in ad hoc experiments. It would also be interesting to understand how and when during development nPAPs are established. Furthermore, as suggested by previous literature ^61^, it would also be important to explore how pathological conditions, such as Parkinson’s disease, may alter the functionality of these patterns and their interaction with cortical regions. Altogether, our results represent a proof-of-principle demonstration that inter-structure communication (in this case, M1–VL/VM) occurs based on hardwired preconfigured dynamics and perhaps also preconfigured anatomical connections, as reported elsewhere ^62^. Deciphering these rules of communication throughout the cortico-BG-thalamic loops would be instrumental in understanding macro-circuit contributions to learning and execution of movements.

## Methods

All experiments were approved by the Animal Ethics Committee of the Institute of Neurobiology at the National Autonomous University of Mexico (UNAM) and conformed to the principles outlined in the Guide for the Care and Use of Laboratory Animals (National Institute of Health). Every precaution was taken to minimize suffering and the number of animals used in the experiments.

### Animals

A total of 42 male mice were used in this study. From those, 31 animals were wild-type C57BL/6 and 11 were VGAT-Cre. Animals were 3-to 5-month-old male mice and were housed in individual acrylic boxes under temperature-controlled conditions with 12 hour light/dark cycles and free access to food and water. Mice used for behavioral protocols (13 C57BL/6 and 4 VGAT-Cre) were water-restricted and consumed their requirements during training sessions (1 to 3 ml in 40 minutes per day). Animals were trained for six days with 24-hour free access to water per week. All experiments were conducted in the light phase of the cycle.

### Surgical procedures

All surgeries were conducted under aseptic conditions. Anesthesia was induced with a xylazine/ketamine cocktail (5/40 mg/kg) and maintained with sevoflurane (0.5-1.5%). A single subcutaneous dose of atropine was applied 10 min before surgery (0.025 mg/mg). The temperature and respiration were constantly monitored during surgery. After the surgical procedures, animals were constantly monitored and administered antibiotics (gentamicine) and analgesics (meloxicam). *Stereotaxic viral infections.* All stereotaxic coordinates were calculated based on the Paxinos and Franklin mouse brain atlas and are reported in millimeters with respect to bregma. All injections were performed at a fixed rate (50 nl per minute) with a Hamilton micro syringe (Neuro-Syringe 7001, 1μL) and an infusion pump (3WPI-UMP3). The viruses AAV5/CamKIIa-hCHR2(H134R)-mCherry-WPRE-PA or AAV5/CamKIIa-hCHR2(H134R)-eYFP-WPRE-PA (purchased from VECTOR CORE, University of North Carolina) were injected unilaterally into two depths of M1 (AP: + 1.34mm ML: - 1.75 mm DV: 1.25 and 1.75 mm) with a final volume of 1000 nl (500 nl in each deep). In VGAT-Cre mice, AAV5/EF1a-DIO-hCHR2(H134R)-mCherry, AAV5/EF1a-DIO-hCHR2(H134R)-eYFP or rAAV5/EF1a-DIO-eArch3.0-eYFP (from VECTOR CORE, University of North Carolina), was injected into the SNr (AP: -3.52 mm ML: +1.5 mm DV: -4.0 mm) with a final volume of 400 nl. *Fixation headpost implants.* Customized titanium headposts were implanted into the skulls of the animals parallel to the midline and fixed with dental cement and two screws placed on the left parietal bone that also served as ground and reference for the electrophysiological recordings (Figure 1A). A metabond-based (C&B Parkell) clear-skull cap was built on the right hemisphere and the craniotomy and fiber optic coordinates were marked. Craniotomies for silicon probe recordings were performed 24 h prior the first recording session and protected with Kwik-Cast (WPI). Each animal was recorded in three to four sessions, one session per day. *Fiber implantation.* For optogenetic manipulation of the SNr, the same headpost implantation procedure described above was performed, but an additional 4 mm fiber optic was directed to the SNr and fixed with dental cement.

### Optogenetic stimulation

Optic fibers (200 µm, diameter) for M1 and SNr stimulation were located in the following coordinates for M1: AP = 1.34 mm, ML = - 1.75 mm, DV = between -0.3 mm and 1.0 mm; for SNr: AP = -3.52 mm, ML = 1.5mm, DV: between -3.9 mm and -4.3 mm. For channel rhodopsin 2 (Chr2) excitation, we used blue light (465 nm) with a maximum power of 19.9 mW. In three animals, we tested different stimulation intensities ranging from 0.3 to 19.9 mW. For optogenetic stimulation of M1 we used four different train stimulation protocols. Trains consisted of five light pulses of 5 ms with 300 ms (3.3 Hz, protocol 1) or 500 ms (2 Hz, protocol 2). Protocols 3 and 4 were identical to protocols 1 and 2 but paired with a single continuous 2100 ms or 3400 ms (300 ms and 500 ms ISI, respectively) stimulus in the SNr (excitation or inhibition depending on the opsin). SNr stimulation started 100 ms before the first stimulus on M1 and covered the whole length of the train. *Stimulation during behavioral execution.* SNr optogenetic stimulation was delivered in 50% of the trials randomly selected. During stimulated trials, stimulation was configured as a closed loop triggered by the minimum displacement of the lever and terminated with the absence of movement or reward delivery.

### Behavioral protocol

#### Apparatus

Animals were trained and recorded in a head-fixed configuration in a customized stereotaxic-based behavioral set. Head-fixed animals rested on an acrylic platform with access to a holding pole and a movable lever located directly under their forepaws (Figure 4A). The lever and holding pole were located 1.2 cm away from the platform. The lever was located under the left forelimb (contralateral to the recording and stimulation sites) and connected to a voltage transductor (1.2 cm = 2.5 mV). All the voltage signals were digitalized and stored at 250 Hz through National Instruments cards. Water rewards were delivered through a water port connected to a solenoid valve. Rewards were signaled with a green LED located 10 cm in front of the animals. Two LEDs located 10 cm to the side of the head of the animal (one to the left, one to the right) were used to specify the duration of the required movement: blue light for long trials (500 ms) and white light for short trials (200 ms). All behavioral parameters were controlled and recorded with customized software programmed in LabView.

#### Task

Mice were trained in a two-interval behavioral protocol. Animals were asked to displace the left lever (push or pull) for at least 1.2 cm in any direction and for at least 200 ms (short trials, indicated by a white LED) or 500 ms (long trials, indicated by a blue LED). Trials were self-initiated and presented in alternating blocks of 20 short or long trials. On each trial, once the spatiotemporal rule was achieved, the green LED indicated the correct trial, and the subject received a drop of water. After being rewarded, animals were asked to stop displacing the lever for at least 1.5 s before a new trial began. Sessions lasted 40 min and mice were trained in one session per day.

#### Training

Animals were accustomed to head fixation and general conditions during four habituation sessions where head fixation time was progressively increased from 5 to 40 min. After habituation, water restriction was started, and mice were progressively trained to displace the lever (from 50 ms to 500 ms) and associate the movement with reward (∼10 sessions; Modeling phase). After the modeling phase, formal training in the two-interval protocol started. All subjects trained for at least 50 sessions before electrophysiological recordings or optogenetic manipulations were performed.

### Electrophysiological recordings

#### General procedures

Recordings were performed using silicon probe microelectrode arrays with a tetrode configuration (NeuroNexus; A4X4-tet-5mm-150-200-121). Craniotomies for recordings were located above the VL/VM complex (1.5×1.5 mm, centered at AP - 1.34 mm, ML + 1.0 mm) and protected with Kwik-Cast (WPI). On recording day, Kwik-Cast was removed, and the array was impregnated with DiI for histological localization of the recording sites. Recording electrodes were inserted in the middle of the craniotomy with slight variations in the antero-posterior and mediolateral axis to only avoid blood vessels. In different recording days, the probe was inserted in the same area. The electrodes were slowly lowered under sevoflurane anesthesia until the desired depth (DV ∼ 4.00 mm). Then, the animals were awakened and allowed to recover for at least 30 min before recordings were started. Recordings in subsequent days were performed through the same craniotomy. After each session, the craniotomy was again protected with Kwik-Cast. To prevent infections and maintain the welfare of the animals, recordings were limited to four sessions per animal.

#### Electrophysiological data acquisition and processing

Wide-band (0.1 to 8000 Hz) neurophysiological signals were amplified 1000 times via Intan RHD2000 series Amplifier System and continuously acquired at 20 kHz. Data visualization and processing were performed from raw data using Neuroscope and NDManager (http://neurosuite.sourceforge.net). Spike sorting was performed semiautomatically using the clustering software KlustaKwik (http://klustakwik.sourceforge.net) and the graphical spike-sorting application Klusters (http://klusters.sourceforge.net) ^63,64^.

### Analysis of neural data

Most of the recordings for M1-evoked thalamic patterns of activity consisted of 50 stimulation trains, which were given every 5 s (sometimes, 100 trains were given; the average protocol duration was 4 - 8 minutes). Neurons were discarded when presented no spiking activity or very low firing rates (< 0.1 Hz) during 5 s (for M1-stimulation experiments) or 8 s (for behavior-related recordings) time windows around the onset of stimulation trains (or reward onset) in more than 40% of the stimulation trains. Due to recording stability conditions, for example, subtle silicon probe movements due to strong animal movements or the natural re-accommodation of the tissue, some neurons were possible to record only in one stimulation protocol or behavioral condition. These neurons were excluded from specific analysis when indicated in the main text. Firing rates were calculated based on inter-spike intervals over the entire recording period. Response latencies for the different components of the response (i.e., short- and long-latency increases and decreases) were calculated by binarizing neuronal data to 1 ms resolution and constructing perievent histograms (-1 to +4 s around the onset of the first stimulus of the train). Then, for each cell, based on the 1 s prior baseline activity, we determined a confidence interval with an upper and lower limit of 99% (for increased responses) and 1% (for decreased responses), respectively. Responses to M1 stimulation were considered significant if they exceeded these limits by at least 1 ms, and the time of the first exceeding bin was defined as the response latency. *Surrogated spike trains.* To construct random spike trains for the random distributions depicted in figures 2b and 5e, for each cell, each spike time from the spike trains obtained from the -1 to +4 s (or -4 to +4) s around the onset of the stimulation (or reward delivery) time windows, were circularly and randomly re-ordered by adding time in a range of ± 1 to 5 (or ± 1 to 8) s. Spike times offsetting outside the intervals were re-accommodated at the beginning or the end of the interval depending on the magnitude and the sign of the offset. This procedure was repeated 100 times. *M1-evoked patterns.* We used the principal component analysis (PCA)/Silhouette-based method reported in ^11,33,65^. In brief, we applied PCA to the z-scored M1 stimulation-evoked patterns of each VL/VM cell in a time window of 1.5 s or 2.5 s for the 300 ms or 500 ms ISI, respectively. To assign cells to specific clusters with similar characteristics, we applied k-means to the first three principal components. To obtain the best classification, we repeated the process 1000 times with projections ranging from 2 to 10 clusters. We scored each projection with the Silhouette method and selected the projections with the highest Silhouettes scores. This procedure detects the general shape of the patterns. To compare between pattern shapes (pattern clusters) evoked by the 300 ms and 500 ms ISIs (Figure 3A), we focused only on the first 300 ms after each stimulus of the train for both conditions; that is, we cut the last 200 ms of the interval for the 500 ms condition. Then we pooled the activity of all recorded cells in both conditions and ran the procedure described above. The same classification method was applied for M1-evoked patterns from trained animals (Figure 6B) and the experiments with paired SNr stimulation (Figure 7G), except that in this case we compared the 300 ms condition from naïve vs expert animals, or the same animal but with or without SNr stimulation, respectively. Hence, in these cases, it was not necessary to segment responses. We also used this classification method for the neural patterns associated with behavioral performance in the two-interval protocol (Figure 5E) between the 200 ms and 500 ms movements. In this case, the temporal domain was normalized to a fixed number of bins (2300). To this end, the 200 ms immediately prior to reward delivery for the 200 ms ISI condition was interpolated to 500 bins. This manipulation equalized the number of bins to 500 ms (500 bins) immediately prior to reward delivery in the 500 ms ISI condition. We complemented both conditions with 1000 ms (1000 bins) before the 200 ms (normalized to 500 bins) and 500 ms (500 bins) windows and 800 ms (800 bins) immediately after reward delivery, for a total of 2300 bins. *Euclidean distance analysis.* To compare population dynamics between 300 ms and 500 ms ISIs (Figure 2) or between stimulation protocols with and without SNr-paired stimulation (Figure 7), we calculated the Euclidean distances from population profiles obtained from the five stimuli’s average M1-evoked responses. Euclidean distance analysis was based on the sorted activity 30-bin matrices in Figure S1 and Figure 7E. In those matrices we calculated the Euclidean distance between each possible pair of bins between two conditions (300 ISI vs 500 ISI). We report the average Euclidean distance and the diagonal asymmetry index. For the diagonal asymmetry index, we first calculated two vectors: one with the distance values at the minimum (minimum distance vector) and a second with the bin difference between the identity diagonal and the minimum values (called the diagonal asymmetry vector). The diagonal asymmetry vector was transformed into angular differences, where a difference of 30 bins between the diagonal and the minimum value corresponds to 360 degrees. For the self-sorted neural sequences where a condition was compared with itself (e.g., 300 ms ISI vs 300 ms ISI), the minimum distance and the diagonal asymmetry vectors were constituted by thirty zeros (Figure 3C, first two matrices, with dots falling exactly over the identity diagonal). In contrast, for cross-sorted sequence matrices, the minimum values of the distance matrix can fall in different values between the rows and columns, and both vectors were different from zero, for example, in the relative comparison between the 300 vs 500 ms ISIs (Figure 3C).

#### Statistical analysis

Electrophysiological and behavioral data are presented as median + 25^th^ and 75^th^ percentiles. Statistical comparisons for electrophysiological or behavioral data between groups were performed with Mann-Whitney or Kruskal-Wallis tests as stated in each section. A Bonferroni post hoc test was used for multiple comparisons. Statistical differences were considered significant if P values were < 0.05.

### Histology

At the end of the experiments, mice were euthanized (pentobarbital 100-150 mg/kg) and transcardially perfused with 4% paraformaldehyde (PFA). The brains were collected and processed to confirm electrode, optic fiber, and infection sites.

## Acknowledgments

We thank Ana Inácio for indispensable technical advice and critical reading of this MS. Authors thank the support provided by all the members of Laboratory A-02 from the Institute of Neurobiology, UNAM; Cuautli Pacheco and Martín García for providing invaluable support in animal maintenance and care; Oscar Prospéro for generous donations of indispensable equipment. Anaid Antaramian and Adriana González from Unidad de Proteogenómica, INB. Jessica Gonzalez-Norris for proofreading. Perla González-Pereyra is a doctoral student from Programa de Doctorado en Ciencias Biomédicas, Universidad Nacional Autónoma de México (UNAM) and supported by fellowship 749154 from CONAHCyT-México.

## Author contributions

Conceptualization: PGP, PRO

Methodology: PGP, PRO

Investigation: PGP, MGMM, DIOR, PRO

Data Curation: PGP, PRO

Formal analysis: PGP, PRO

Writing—original draft: PGP, PRO

Writing—review & editing: PGP, PRO, HM, LT

Resources: LT, HM

Supervision: PRO

Project administration: PRO, CIPD

Funding Acquisition: PRO

## Supplementary Figures

**Supplementary Figure 1.**
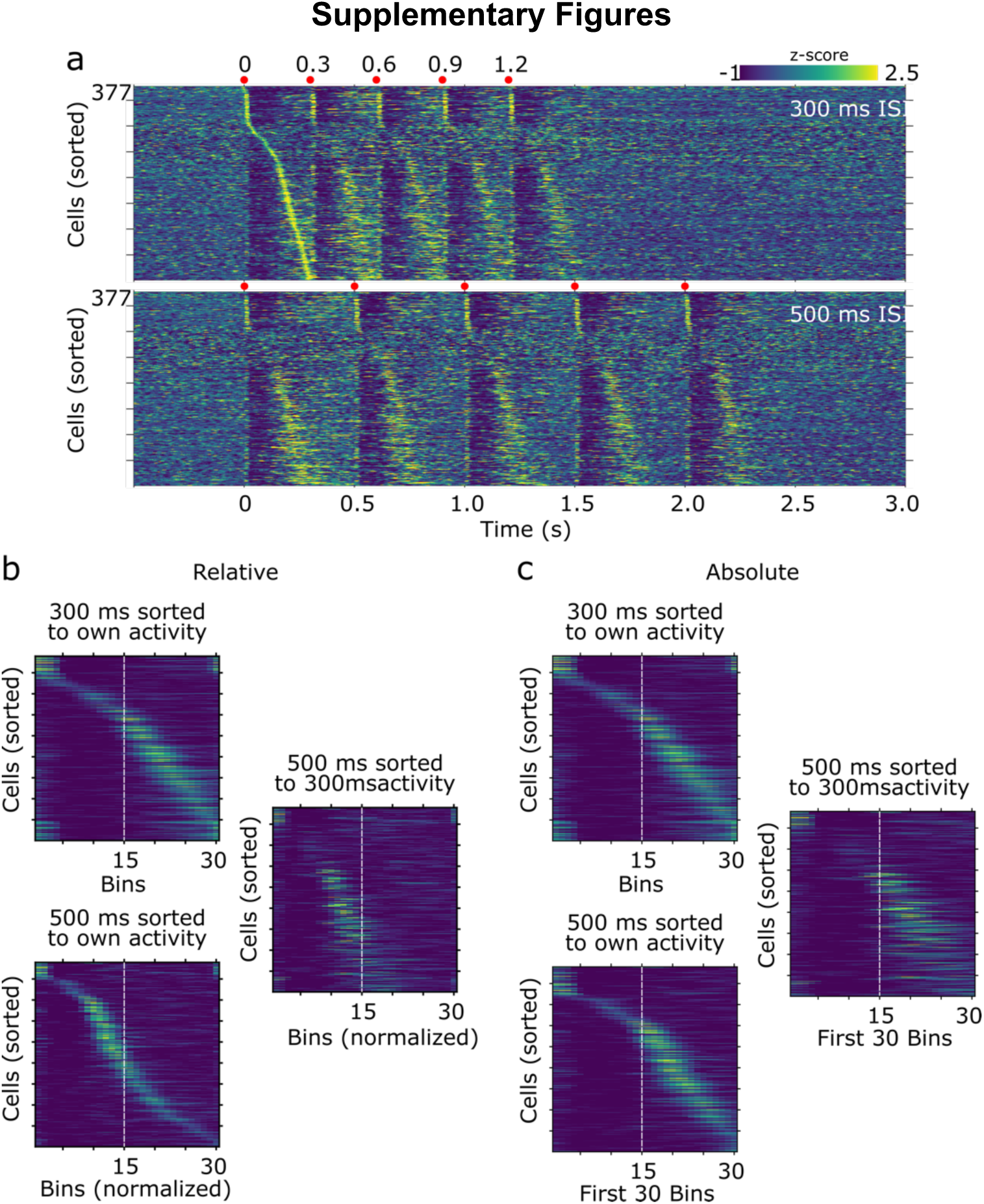
**a**) Average firing rates (z-scored) evoked by M1 stimulation for cells recorded under two conditions: 300 ms (top) and 500 ms (bottom) ISls. Cells in both matrices were sorted according to the moment of their highest firing rate between the first and second stimulus of thee 300 ms ISI train. Each stimulus of the train is indicated on top of each panel (red dots). Thirty-bin matrices depicting the average firing rates for the 5 stimuli of the train for relative (**b**) and absolute (**c**) comparisons. Cell sorting is indicated on top of each panel. For 500 ms relative matrices, neural activity was adjusted to 30 bins. For 500 ms absolute matrices, neural activity corresponds to the first 300 ms after the onset of stimulation. White dashed lines are depicted as visual reference.

**Supplementary Figure 2.**
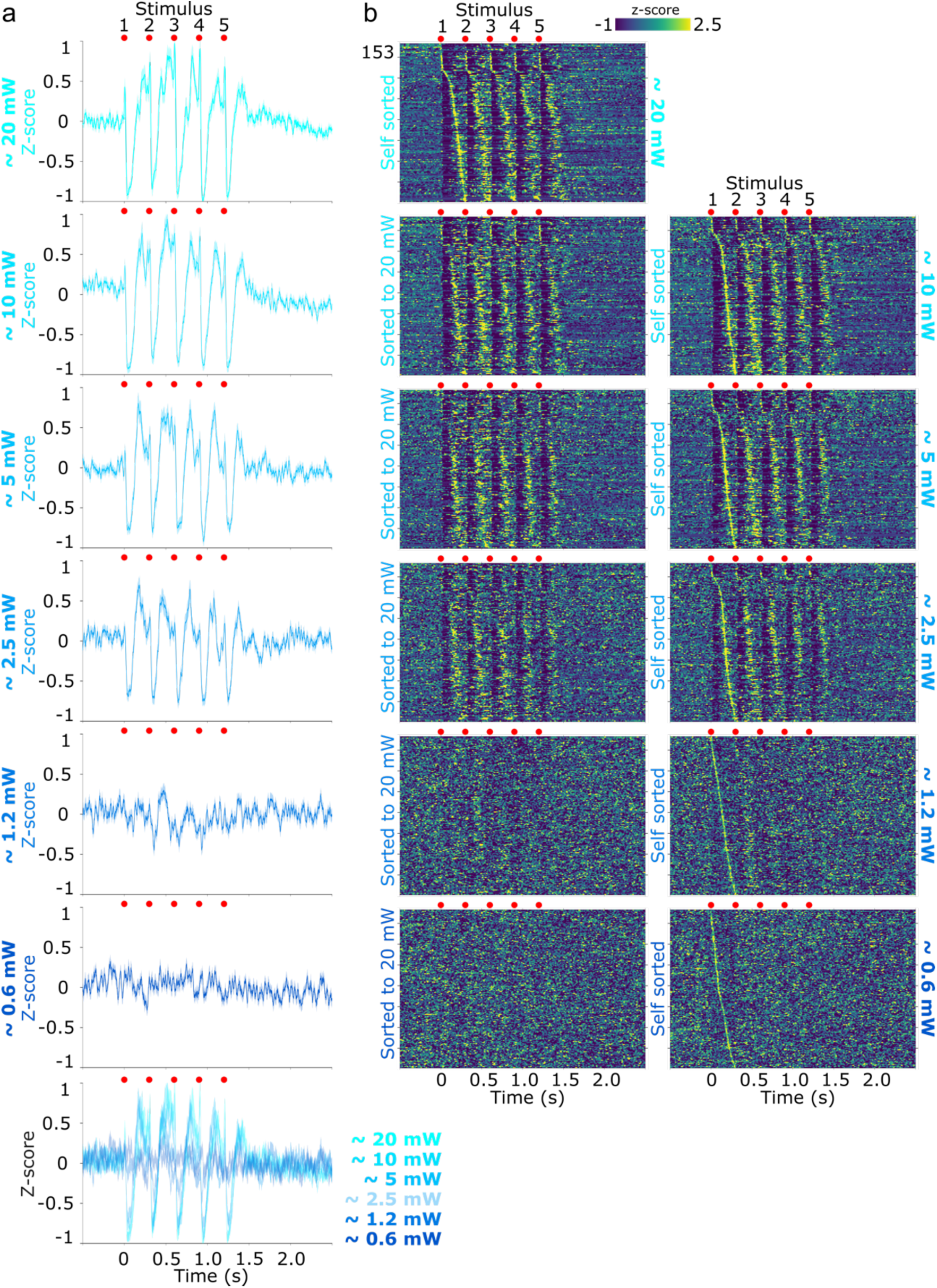
M1-evoked response patterns in urethane anesthetized animals. **a**) Full train-averaged population response aligned to the first stimulus of the train for six light intensity stimulations (color code indicated on the Y axis) in naïve awake animals (same color code for the rest of the figure). For visual comparison, the bottom panel merged all conditions. **b)** Average firing rates (z-scored) evoked by M1 stimulation for the same cells recorded under the six light intensities. In the left column, cells were sorted according to the moment of their highest firing rate between the first and second stimulus of the train for the 20 mW stimulation. In the right column cells were sorted according to the moment of their highest firing rate between the first and second stimulus of the train for each intensity (self-sorted). Fifty 300 ms ISI trains were given for each light intensity. Light intensities were provided in descending order starting by 20 mW.

**Supplementary Figure 3.**
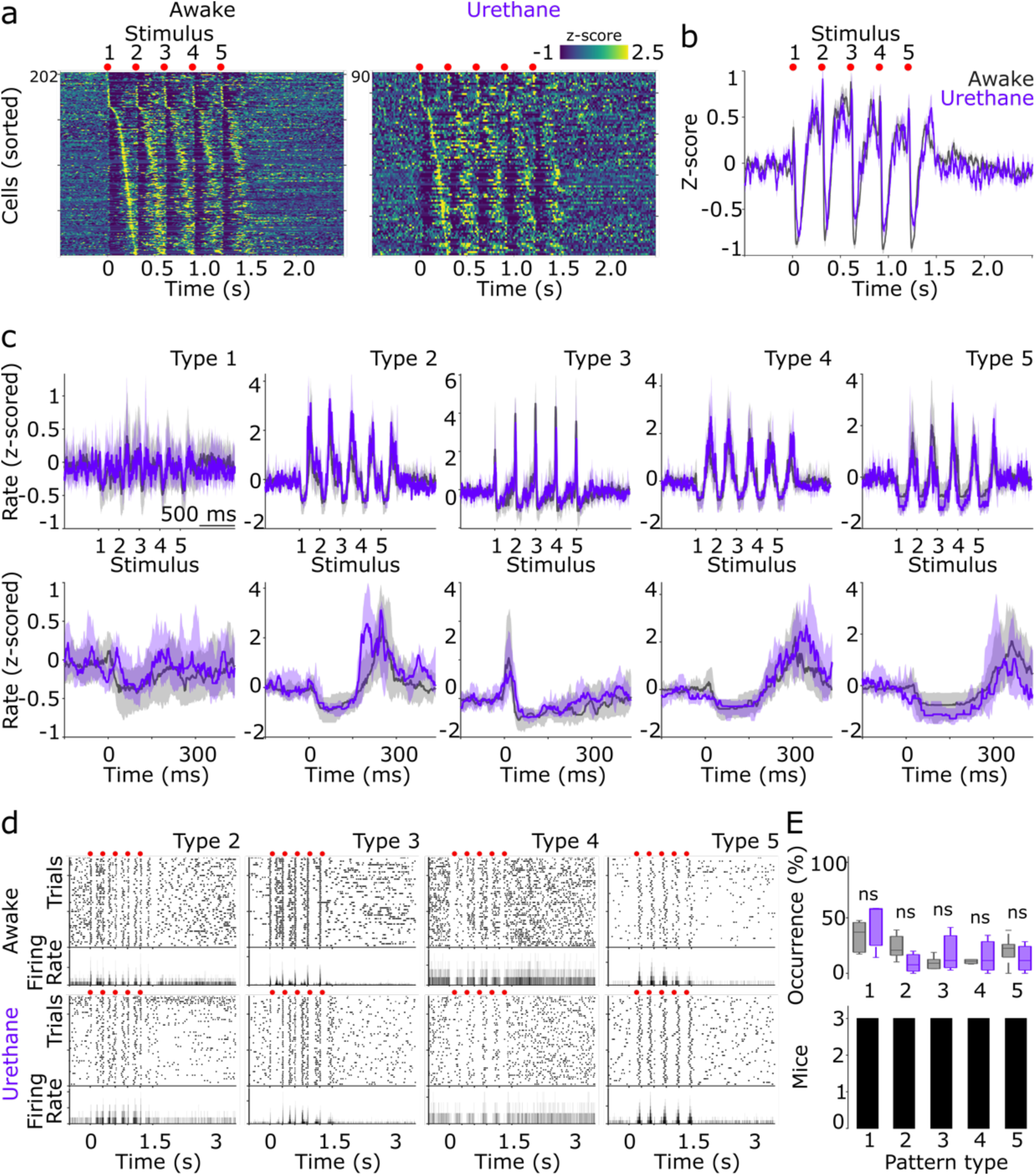
M1-evoked response patterns in urethane anesthetized animals. **a)** Average firing rates (z-scored) evoked by M1 stimulation for cells recorded under two conditions: 300 ms ISI awake (left) and 300 ms anesthetized (right) inter-stimulus intervals (ISIs). Cells were sorted according to the moment of their highest firing rate between the first and second stimulus of the train. Each stimulus of the train is indicated on top of each panel (red dots; same for the rest of the panels). **b**) Full train-averaged population response aligned to the first stimulus of the train for the awake (gray traces) and anesthetized (purple traces) conditions (same color code for the rest of the figure). **c)** Average M1-evoked patterns for cells classified as part of specific pattern clusters for the awake and anesthetized conditions. Solid lines and shaded areas represent the median and the 25^th^ and 75^th^ percentiles, respectively. **d**) Representative spike rasters and their corresponding average peri-event histograms for eight different neurons belonging to specific pattern clusters recorded in awake (top row) and anesthetized (bottom row) conditions. Activity was aligned to the first stimulus of the train. **E**) Percentage of cells belonging to each pattern cluster displayed in A (upper panel). Number of animals that presented each pattern (lower panel). Boxplots indicate median and 75^th^ and 25^th^ percentiles. Statistical comparisons were performed by applying Kruskal-Wallis.

**Supplementary Figure 4.**
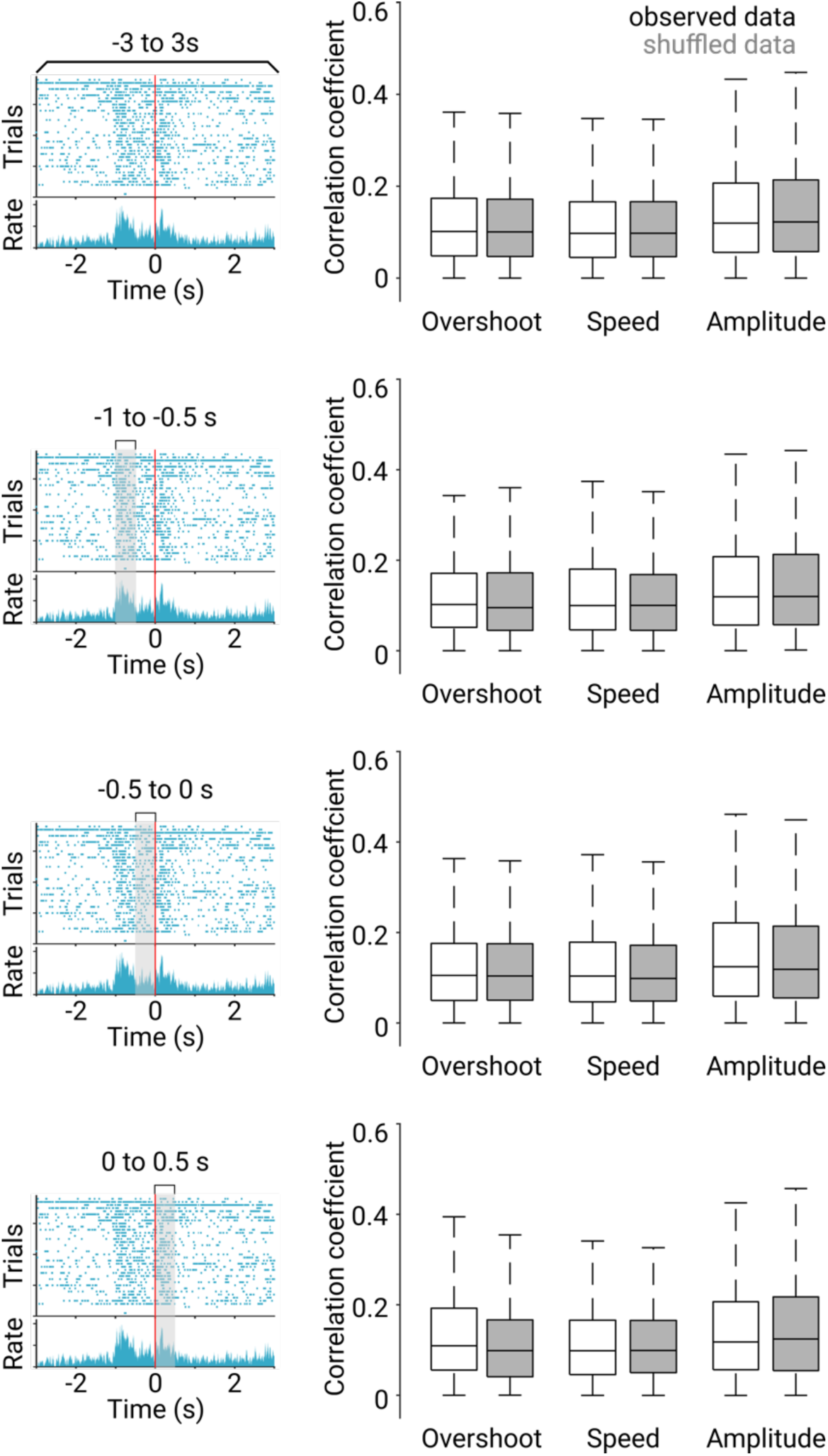
Correlation coefficients (absolute values) for all recorded neurons and behavioral variables. Correlations were calculated between each variable and the average spiking activity from the periods indicated on top of the left rasters (displayed for reference). The coefficients were obtained from the actual data (with boxplots) and from surrogated spike trains of the same data but with randomly shuffled spiking activity (gray boxplots). Boxplots indicate median and 75^th^ and 25^th^ percentiles. Statistical comparisons were performed by applying Kruskal-Wallis and Bonferroni post hoc tests.

**Supplementary Figure 5.**
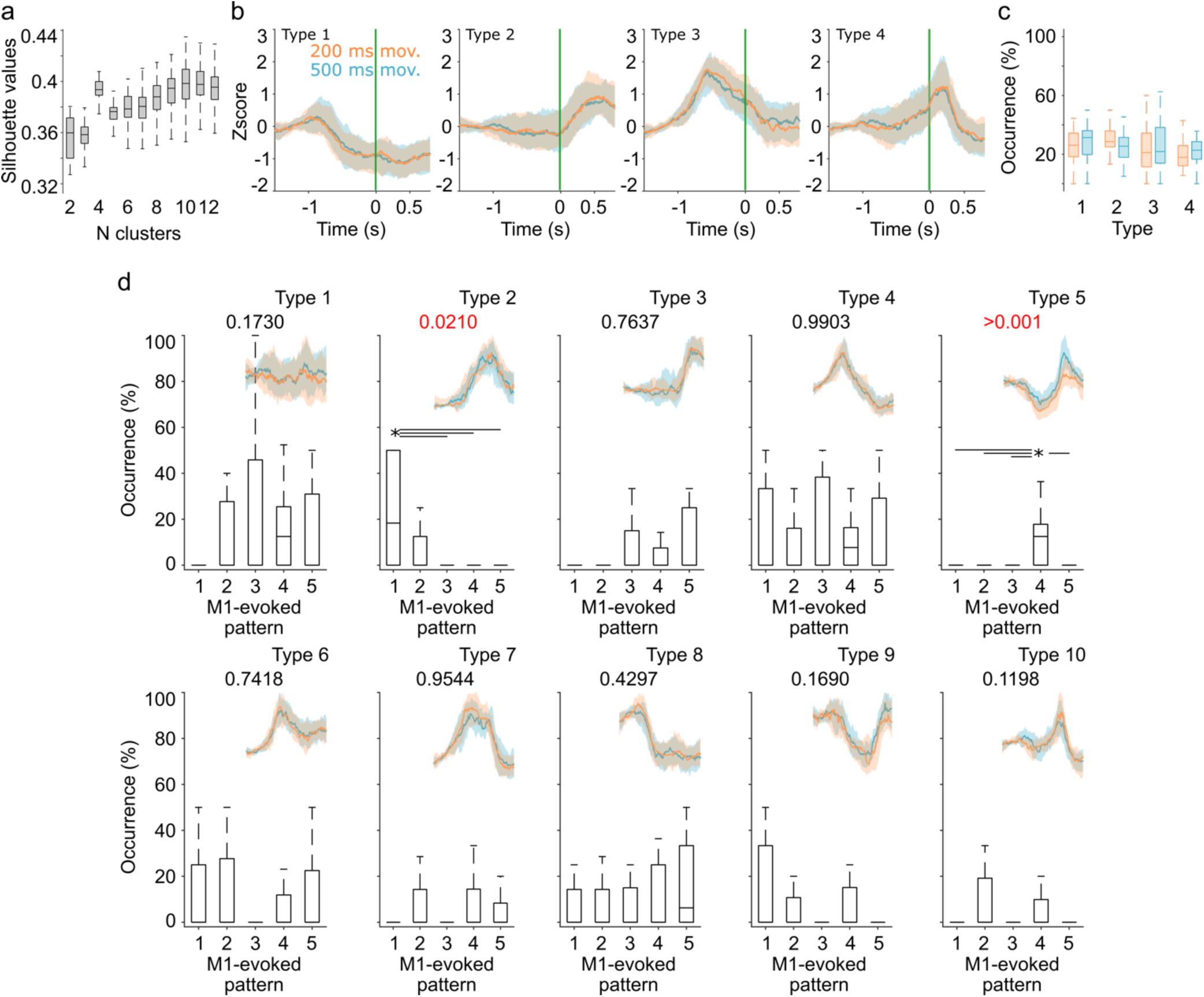
**a**) Silhouette values for 1000 iterations in 2-12 k-means projections from the PCA on the perievent histograms of the spiking activity during movement execution. **b**) Average behaviorally evoked perievent histograms for cells classified as part of specific pattern clusters for the best four-cluster projection for the 200 ms (orange) and 500 ms (blue) conditions. **c**) Percentage of cells belonging to each pattern. Percentages were calculated using sessions with more than 10 cells simultaneously recorded. **d**) Percentage of cells belonging to each M1-evoked pattern (displayed in Figure 6B) as a function of each of the 10 behaviorally associated patterns (displayed as insets on each panel and taken from Figure 5E). Percentages were calculated using sessions with more than 10 cells simultaneously recorded. Boxplots indicate the median and 75^th^ and 25^th^ percentiles. Statistical differences are indicated by asterisks and lines joining specific comparisons (Kruskal-Wallis indicated on top of each panel and Bonferroni post hoc test, P < 0.05).

**Supplementary Figure 6.**
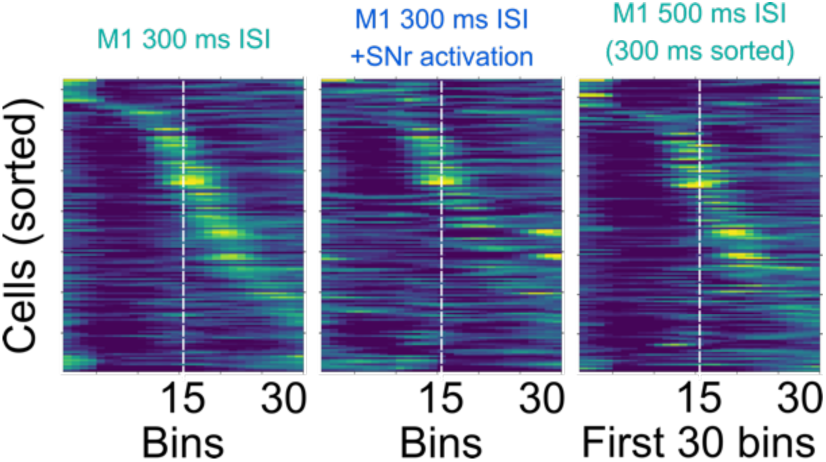
M1-evoked sequential activation matrices (30 bins, 5 stimuli averaged) for 300 ms ISI trains with (middle) and without (left) SNr paired stimulation and for 500 ms ISI (right). The same neurons are presented in the three matrices and were sorted according to the moment of their highest firing rate during the 300 ms ISI (left).

